# Lupus disease flares are concordant with immune responses to blooms of lipoglycan-expressing *Ruminococcus blautia gnavus* strains arising from unstable gut microbiota communities

**DOI:** 10.1101/2022.06.28.498027

**Authors:** Doua F. Azzouz, Ze Chen, Peter Izmirly, Lea Ann Chen, Zhi Li, Alejandro Pironti, Gregory G. Putzel, Chongda Zhang, Adriana Heguy, Mala Masson, Dominik Schwudke, David Fenyo, Nicolas Gisch, Jill Buyon, Alexander V. Alekseyenko, Gregg J. Silverman

## Abstract

Whereas genetic susceptibility for Systemic Lupus Erythematosus has been well explored, the precipitants for clinical disease flares remain a mystery. To investigate for dynamic-relationships between gut-microbial communities and Lupus disease activity, we performed taxonomic surveys of fecal 16S rRNA gene amplicon-libraries from Lupus patients and healthy volunteers, obtained at serial timepoints over many months to several years. Individual Lupus patients commonly displayed imbalances in alpha and beta microbiota-diversity, which were uniquely different from healthy individuals as well as from other Lupus patients. Moreover, multivariate analysis of sequential Lupus libraries documented community-wide ecological microbiota instability overtime, most pronounced in patients with Lupus Nephritis LN), a severe form associated with worse prognosis. Lupus gut communities displayed transient spikes of pathogenic bacterial species, with by far the most prevalent being blooms of *Ruminococcus blautia gnavus* (RG), occurring in nearly half of LN patients often concordant with disease activity flares. RG strains isolated during disease flares, but not those isolated from healthy individuals or patients with inflammatory bowel disease, commonly expressed a novel, highly immunogenic cell wall-associated lipoglycan with conserved structural features that include a diacyl glycerol anchor. Cross-reactive antigenic determinants on these lipoglycans were recognized by murine monoclonal antibodies, and by spontaneously arising Lupus serum IgG antibodies with peak serum antibody responses also concordant with RG blooms. As SLE is frequently characterized by remitting-relapsing disease, despite appropriate treatment, we speculate that gut blooms of pathogenic bacteria, that impair gut barrier function and stoke systemic inflammation, directly contribute to immunopathogenesis.

## Introduction

The interconnected multidimensional commensal microbial communities that we harbor within us provide a myriad of functions critical for both nutrition and general health. Our commensals also serve as early sparring partners, essential for the development and maintenance of the most fundamental layers of cellular defenders from the innate and adaptive immune systems. In health, there are complex relationships within these microbiota communities that are believed to maintain and reinforce a dynamic equilibrium. There is however, mounting evidence that imbalances, termed dysbiosis, within gut commensal communities are common in patients with diverse inflammatory and autoimmune diseases (reviewed in ^1^). Such imbalances are postulated to unleash the pathogenic potential of inherited susceptibility factors that otherwise remain dormant in unaffected relatives.

Lupus is the archetypic systemic autoimmune disease that carries great morbidity and mortality. Much is known about the immunologic pathways associated with pathologic autoimmunity and tissue injury. Yet despite progressive advances in the therapeutic armamentarium and even with regular clinical evaluations, many patients nonetheless suffer from inadequate clinical responses and recurrent flares of disease ^2^. Amongst the greatest morbidity is that incurred by renal involvement that affects up to 60% of patients, with recent reports estimating that 20% will continue to progress to end-stage renal disease (discussed in ^3^).

In our earlier cross-sectional studies of the gut microbiomes of Lupus patients, we characterized the communities within an urban female cohort with great heterogeneity of clinical features and organ system involvement ^4^. From these cross-sectional studies, direct correlations were documented between SLE disease activity index (SLEDAI) and dysbiotic shifts associated with reduced community diversity and richness. Many clinically active patients also had gut expansions of *Ruminococcus gnavus* (RG), a species reassigned to the genus Blautia in the Lachnospiraceae family of spore-forming obligate anaerobes of the *Clostridium* cluster XIVa of the phylum Firmicutes ^5^, which is one of the 50 most commonly found species in the intestine albeit with low abundance levels of ∼0.1% in healthy adults ^6^. Notably, in a large untreated Lupus cohort in China, RG was also recently described as “the most discriminative species enriched in the gut microbiota associated with Lupus Nephritis (LN) ^7^. This suggests broader relevance of RG expansions to Lupus pathogenesis across diverse epidemiologic populations, ethnicities, and geography ^7^. Yet, there are few instances in which individual gut species have previously been implicated in clinical autoimmune pathogenesis. We have hypothesized that both quantitative changes (i.e., blooms), as well as genomic and phenotypic differences in RG strains, may contribute to Lupus pathogenesis.

Within the current studies, we have investigated the temporal dynamic stability of the Lupus gut microbiota community. For this purpose, we assembled a collection of fecal samples from Lupus patients and healthy control (CTL) subjects spanning from several months to years (see Methods and Supplementary Table 1), which were used to newly generate a series of 16S rRNA amplicon libraries. Remarkably, we found that Lupus patients commonly display gut microbiota community instability. Overtime, within these distorted communities dramatic but ephemeral blooms of several bacterial pathogens were documented. Most notably, gut spikes in the abundance of RG, an obligate anaerobe, arose concordant with significant renal flares in nearly half of LN patients.

To characterize the features of strains responsible for RG blooms, individual RG colonies were isolated from the fecal samples of two patients at the time of their LN flare, and these RG strains were found to produce a novel immunogenic lipoglycan, composed of a diacyl-glycerol lipid anchor and associated saccharides, and purified lipoglycan from different RG strains shared common structural and common antigenic features. Notably, these lipoglycan molecules are physically associated with the *R. gnavus* bacteria represented in blooms occurring during (or that directly cause) disease flares.

Such RG lipoglycans are recognized by IgG antibodies secreted into the circulation of LN patients. We hypothesize these antibodies are integral to the immune complexes that cause Lupus disease, tissue damage and especially in glomerular injury (i.e., immune-mediated renal damage that can result in renal failure). These findings provide the first evidence of previously unsuspected features of the intimate relationship between intestinal microbiota dynamic instability, blooms of a pathobiont, and Lupus pathogenesis.

## Results

### Gut dysbiosis in Lupus patients with high disease activity and active renal involvement

To study the gut microbial communities in sequential samples from disease-affected individuals, we examined two to six fecal samples from 16 individual patients, representing 44 samples obtained over 24-291 weeks following initial recruitment (Supplementary Tables 1&2). For comparisons, we also evaluated 72 samples from 22 healthy adult female CTL volunteers, which included 2 to 12 samples from 9 individuals. To assess the diversity within these bacterial communities, fecal DNA was extracted, followed by targeted amplification of a standard gene fragment to generate 16S rRNA gene amplicon libraries. These 114 libraries were sequenced in two batches with consistent results in technical duplicates (Supplementary Figure 1). After sequence analysis and data filtering, each library had between 10,345 and 307,479 reads assignable to Amplicon Sequence Variants (ASV) with known taxonomic associations, which enabled estimations of the relative abundance of diverse taxa in each sampled community.

In these analyses, patients who fulfilled ACR criteria for renal involvement at any time ^8^ were here referred to as LN (Supplementary Tables 1A&2A), and these patients were then dichotomized based on renal disease activity at the time of sampling as inactive or active, based on urinary protein creatinine ratio (uPCR) <0.5 or > 0.5 respectively, as defined in the SLEDAI domain. Others are referred to as non-renal patients (Supplementary Tables 1B&2B). Based on earlier microbiome studies ^4^, we designated those with overall SLEDAI score of ≥8 as high disease activity, with others as low disease activity. These gut community findings reiterated our earlier reported results ^4^ that high SLEDAI scores and active LN were both associated with decreased richness/diversity alpha diversity of microbiota communities, reflected in several measures of richness and evenness in estimates of alpha diversity, compared to CTL subjects (Supplementary Figures 2-5).

### Lupus disease activity is associated with greater phylogenetic diversity

Based on Principal Coordinates Analysis (PCoA), the communities recovered from Lupus patients displayed significant differences overtime in beta diversity compared to CTL using Jensen-Shannon divergence (JSD) dissimilarity metric (multivariant distance Welch’s W_d_* test, *p*= 0.001) ^9^(Figure 1A). Furthermore, there were significantly greater differences with communities associated with high disease activity (*p*= 0.001) (Figure 1B), and in communities from patients with active LN (*p*= 0.001)(Figure 1C).

**Figure 1.**
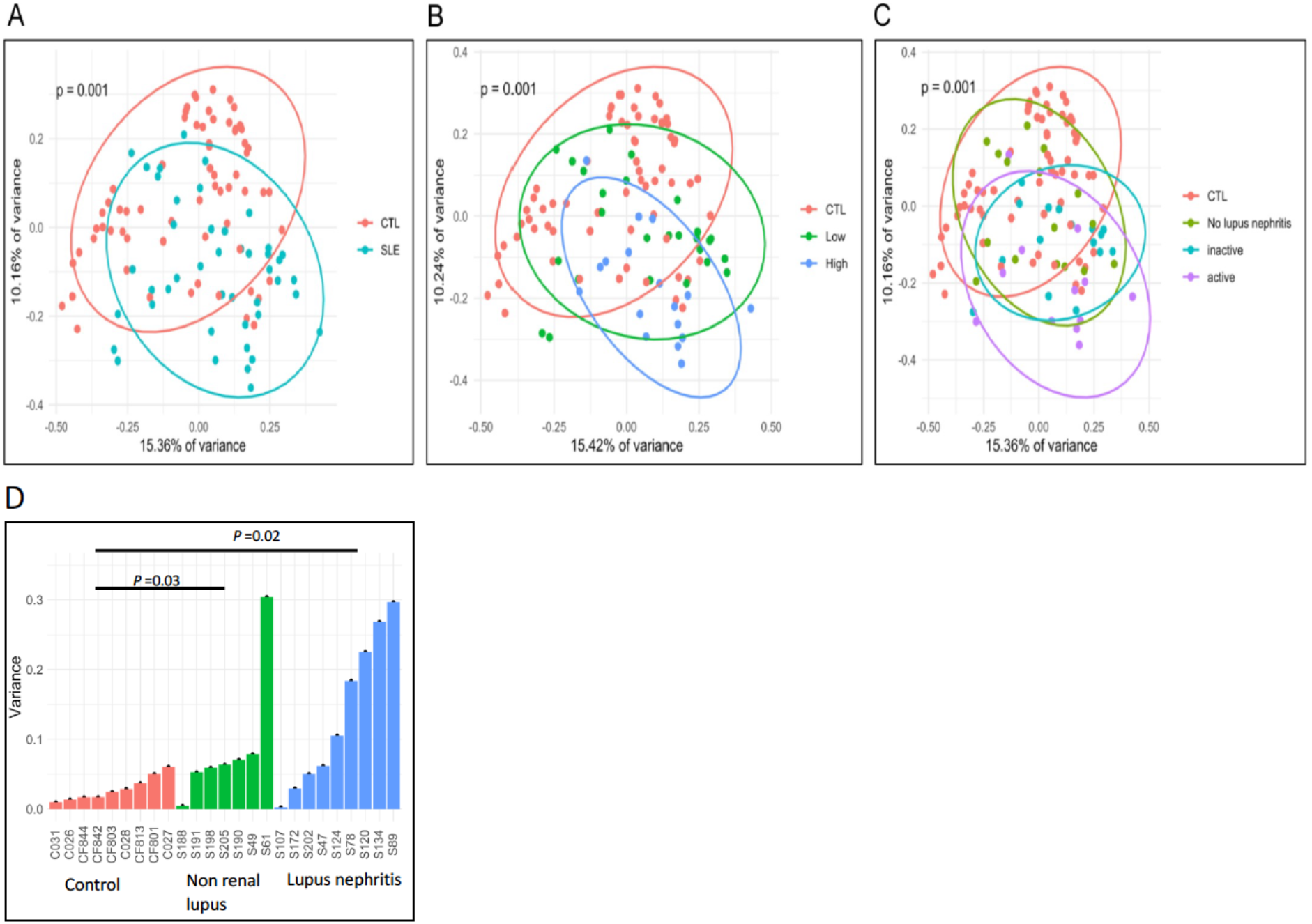
Dysbiosis and longitudinal instability in SLE microbiota communities compared to healthy individuals. A) Principal Coordinates Analysis (PCoA) was used to estimate beta diversity between groups using the Jensen-Shannon divergence (JSD) dissimilarity metric. Commensal communities from SLE patients were heterogenous, with most exhibiting significant distance variance from CTL (multivariate distance Welch W_d_* test, *p*=0.001). B) Compared to CTL, variance in diversity was greatest within the SLEDAI high subgroup, and C) the subset with active renal disease. D) To compare the overall dynamics of shifts in fecal communities sampled overtime in different subjects, subject variances were computed based on JSD using the average multivariate dissimilarity estimation method reported in ^9^. The variances in these three groups were significantly different (Kruskal Wallis ANOVA, *p* = 0.03). Patients with a history of nephritis were assigned to the renal group based on ACR criteria, whereas the patients in the non-renal group were without a history of documented Lupus nephritis. Lupus patients have more unstable gut microbiota than healthy individuals Variance of gut microbiota was significantly different in the renal Lupus group, compared to healthy subjects (two-sided Mann-Whitney test, *p* = 0.02). The non-renal Lupus group was also significantly different than the healthy subjects (*p* = 0.03). Notably, the overall variance in the renal group was not different from the non-renal group (*p* = 0.379, NS).

### Instability overtime is a common feature of Lupus gut microbiota communities

We next asked whether variance within the composition of the libraries from individual Lupus and healthy donors was stable overtime. To assess for stability of gut microbiota overtime community-wide multivariate analyses were performed on the multiple libraries from 16 Lupus-affected individuals and 9 healthy subjects, without assumptions based on time-intervals. Using a proven approach for estimating averages within group dissimilarity ^9^, we found limited differences in overall composition amongst the sampled communities from each of the CTL subjects, while the level of variance was remarkably variable between communities sampled at different timepoints from the same individual Lupus patient (Kruskal Wallis ANOVA, *p*=0.03) (Figure 1D). Notably, compared to controls, there was much greater variance within the Lupus subgroups; non-renal (two-tailed, *p*=0.03), and renal (LN) (*p*=0.02) (Figure 1 D). However, we found no overall correlations between community variance; with disease duration, the periods between sample collection, the maximal disease activity, the range of disease activity in different visits (Supplementary Tables 1&2), or the medication regimen in an individual subject (Supplementary Table 3). Furthermore, the variance in the renal and non-renal groups were not significant different (*p*=0.38, NS). Taken together, these findings suggest the gut microbiota communities in Lupus patients are inherently unstable in microbial composition overtime. These observations suggested that further taxonomic investigations were needed, in part to test the hypothesis that within these unstable community milieus specific bacterial species underwent expansions.

### Dynamic blooms of individual bacterial species are common within Lupus gut microbiota communities

In-depth analyses of the sequential microbiota libraries revealed that in several individual Lupus patients there were transient, but often pronounced blooms of ASV-defined taxa within both Veillonella and Fusobacterium genera, which were absent in all healthy subjects (Supplementary Figures 6&7). Whereas *Veillonella*, a gram-negative Firmicutes genus, can be a normal member of the intestinal or the oral mucosal communities, these can also represent local outgrowths, representing translocations from the oral cavity into the intestine, or even overt infections ^10^. The *Fusobacterium* genus of anaerobic gram-negative non-spore-forming bacteria are generally considered oral pathogens ^11^. Notably, there were *no* associations between these types of blooms with specific Lupus clinical features or organ involvement, and neither temporal associations between Veillonella nor Fusobacterium blooms nor with overall flares of Lupus disease activity (Supplementary Figures 6&7, Supplementary Tables 1 and 2), as shown in taxonomic surveys at a genus and species level (Supplementary Figure 8).

### RG blooms were concordant with Lupus disease flare episodes

In our earlier cross-sectional report, we documented a mean 5-fold over-representation of RG within a larger SLE cohort, compared with the libraries from controls ^4^. In the current studies, we also found quantitative increases in RG relative overall abundance (Wilcoxon, *p*= 0.076) (Supplementary Figure 5A). Importantly, increases in RG abundance were most marked in patients with high disease activity, with SLEDAI scores of 8 or greater (Wilcoxon, *p*= 0.01) (Supplementary Figure 5B), and in those with active renal disease (Wilcoxon, *p*= 0.02) (Supplementary Figures 4&5C)(Supplementary Figure 8). Furthermore, when examined in a continuous distribution, there was a modest but significant direct correlation between disease activity and RG abundance (Spearman, *r* = 0.320, *p*= 0.034). Overall, these findings therefore reaffirmed earlier associations between RG expansions within Lupus microbiota communities, proteinuria, and high disease activity in general.

To investigate for temporal changes in RG abundance in these patients, we closely examined the sequential libraries of the CTL and Lupus patients. These surveys included 49 values from 9 HC, 28 samples from nine LN patients (identified as S47, S78, S89, S107, S120, S124, S134, S172, S202), and 16 values from seven other patients who never had renal manifestations (i.e., non-renal patients; S49, S61, S188, S190, S191, S198 and S205) who instead had histories of inflammatory arthritis, cutaneous disease, and/or other disease features (Supplementary Tables 1A&B). In accord with the level reported in a large-scale population-based survey, we found an average 0.15% RG abundance, with remarkable stability within the healthy female CTLs (Figure 2A). Amongst the 16 Lupus patients, 11 patients exhibited low or undetectable level of RG abundance in their gut communities, with limited variation overtime in individual patients (Figure 2B). Of these 11 patients, for patient S134, who had quiescent LN during our study (Supplementary Table 1A, also had low or undetectable RG in all four timepoints sampled. Similarly, another five LN patients also displayed stable low RG abundance pattern despite episodes of active renal disease documented at one or more of the sampling timepoints (Figure 2B).

**Figure 2.**
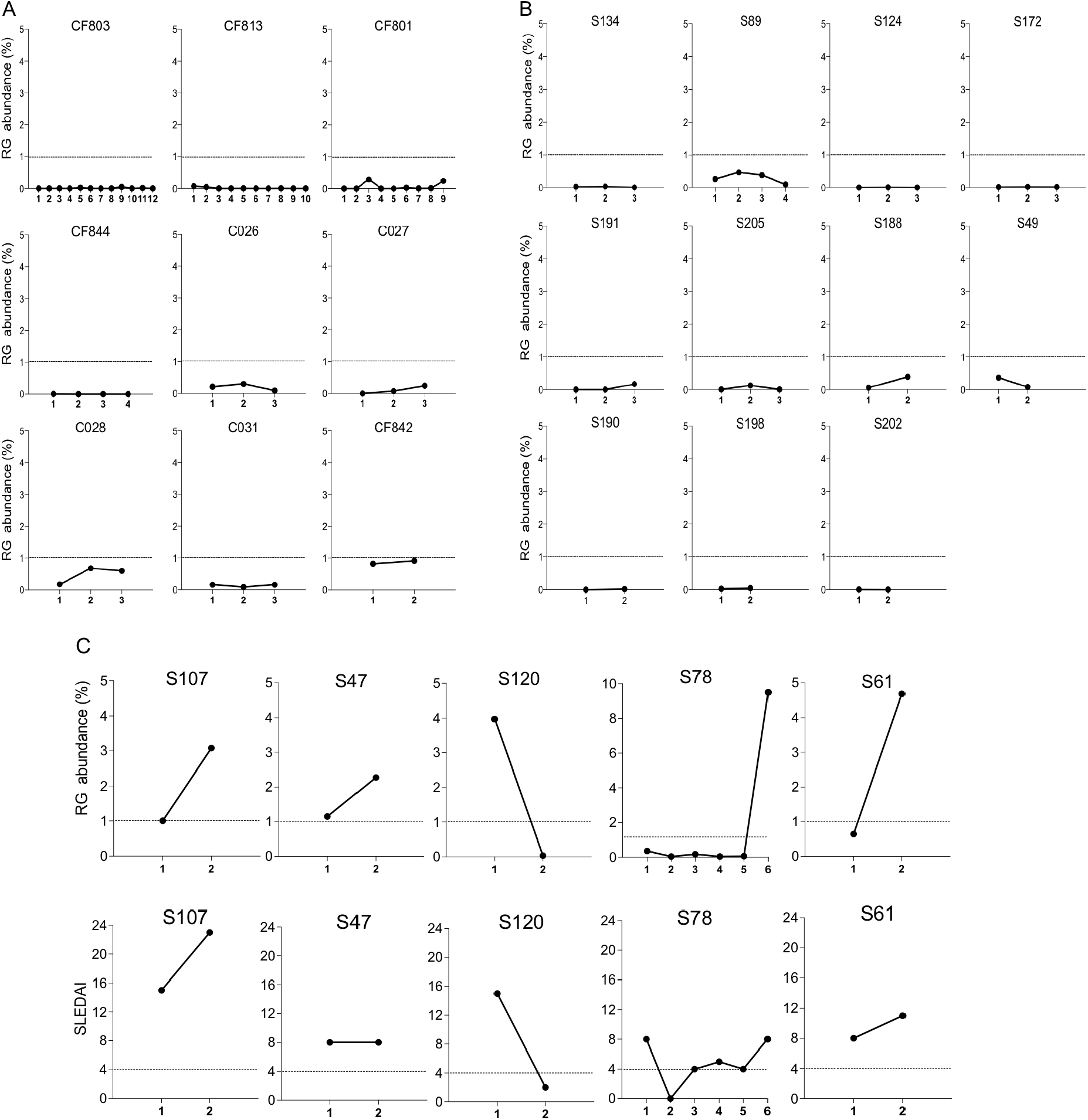
Dynamic changes in RG abundance documented at sequential time points in healthy and Lupus-affected individuals. A) All healthy control subjects displayed a stable low abundance in RG representation. B) In 11 of the 16 SLE patients under investigation, a stable low abundance in RG representation was detected. C) In 5/16 (31%) of the SLE patients evaluated overtime, and the abundance of RG fluctuated greatly overtime. In these cases, RG abundance at much higher levels were present in fecal samples obtained proximal to visits in which disease flares were documented. In all but one of these RG bloom-associated patients had documented LN. RG relative abundance was evaluated for 16 SLE in 44 samples obtained at different time points, and for CTL subjects in 49 samples obtained at different time points, which ranged from 2-12 samples per donor. Clinical and demographic data are shown in Table 1. Dotted line depicts an arbitrary 1% threshold of 16S rRNA amplicon representing RG abundance that is highly above the mean 0.15% level in these healthy controls. Abundance levels above 1% were considered a bloom. Note that in panel C, for patient S78 the greater range of RG abundance necessitated a different scale. SLEDAI ≥ 8 was considered high disease activity.

In contrast, the four other LN patients (∼44%) (S107, S47, S120 and S78), dramatic changes in RG abundance were detected. Each had an RG bloom that reached a mean of 9-fold higher abundance than in other sample(s) from the same donor. Such an RG bloom, with temporal concordance with clinical disease flare, was also identified in patient (S61) who had non renal involvement, with inflammatory arthritis (Figure 2C). Taken together, when only samplings from these 5 patients with an episode of RG bloom were examined, the level of disease activity significantly correlated with the RG abundance (*r*= 0.320, *p*= 0.03). Hence, within individual Lupus patient with an unstable gut communities, RG, which colonize the host intestine, with close proximity to the mucosal epithelial barrier ^12^, underwent blooms temporally concordant with disease flares. We therefore speculate that RG blooms may contribute to the clinical pattern of relapsing-remitting disease activity that is widely documented in many Lupus patients despite close clinical monitoring and treatment ^13^.

### Isolation and characterization of RG strains from LN patients at time of disease flare

To investigate the genomic relatedness of strains that colonize Lupus patients, we characterized the genomes of RG isolates from fecal samples of LN patients. Whereas selective media for RG *in vitro* culture are currently unknown, initial efforts using fecal samples from patients at time of low disease activity were nonproductive. However, with fecal samples from two active LN patients, S47 and S107, obtained at time of LN clinical flares and concurrent RG blooms documented by 16S rRNA amplicon analysis (Figure 2C). Colonies were picked, isolates characterized, and whole genome sequences determined. The genomes of *bona fide* RG isolates were compared to the RG1 and RG2 type strains (Figure 3A). Comparisons to all known RG isolate genomes documented that the S47-18 and S107-48, although from different Lupus donors, displayed a high level of relatedness (Figure 3B) (Supplementary Figure 9). By contrast, despite an earlier report that IBD derived from a separate RG clade ^14^, by these analyses these IBD genomes are in fact dispersed throughout the dendrogram of known RG genomes (Figure 3B) (Supplementary Figures 9).

**Figure 3.**
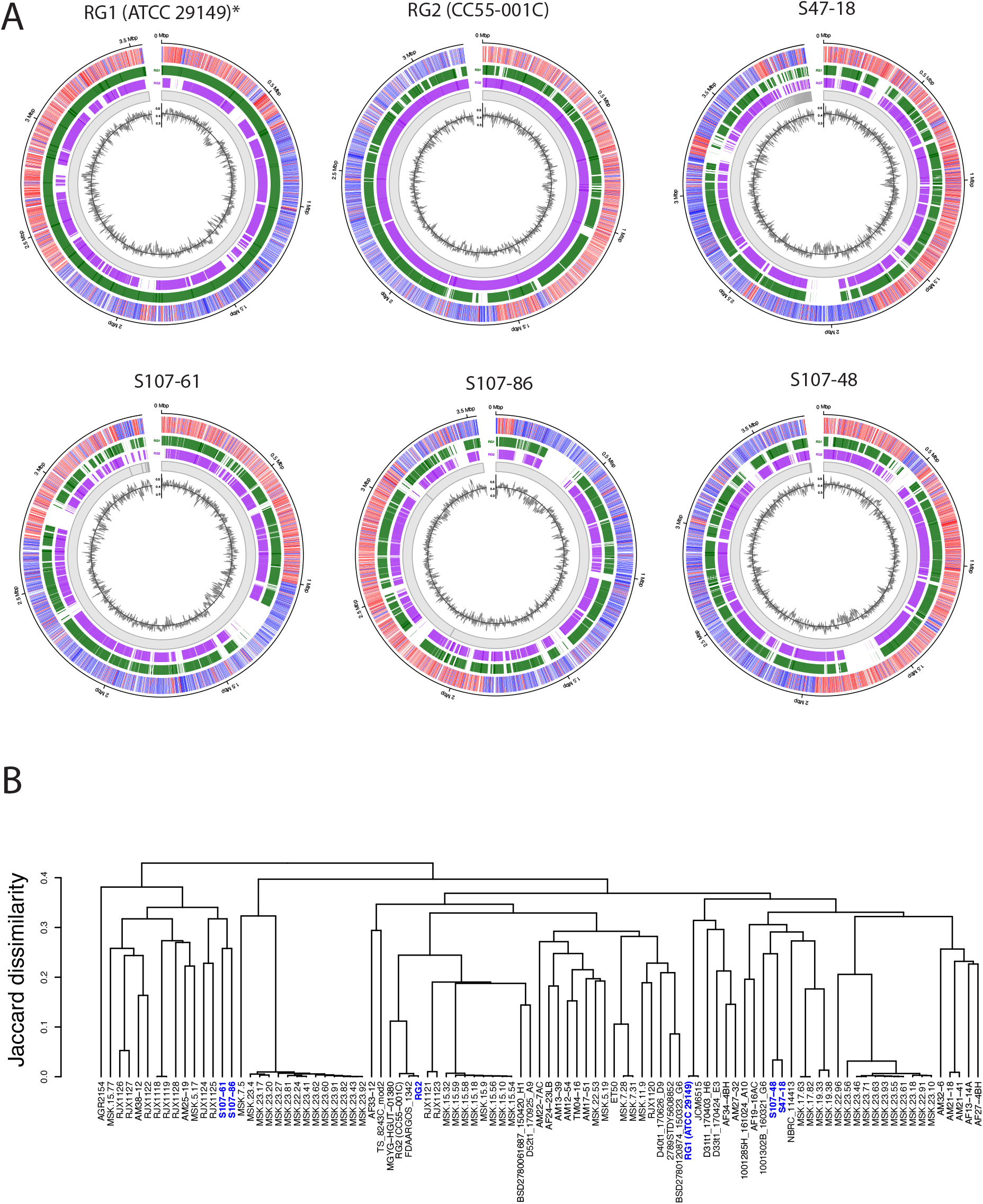
Analyses of *Blautia (Ruminococcus) gnavus* (RG) isolate whole genomes from two Lupus patients in clinical flare. A.) Long-read assemblies of six RG isolates. From inside to outside, circular tracks show average GC content across 1 kbp windows (black line), assembled contigs (gray ring sections), locations of 1kbp windows with BLAST hits to type strain RG1 (ATCC29149) (purple) and RG2 (CC55_001C) (green) genome assemblies downloaded from RefSeq, and locations of genes as predicted and annotated by Prokka colored by strand (+: red, -: blue). The RG1* assembly was downloaded from NCBI, while all others were *de novo*. B) Dendrogram showing complete-linkage hierarchical clustering of the gene content (orthogroup presence/absence) of newly sequenced RG isolates together with assemblies downloaded from NCBI RefSeq. Newly sequenced isolates, together with strain RG1 assembly, are shown in blue.

### LN serum antibodies recognize conserved non-protein oligoband antigens in RG strains from LN

In an earlier report of serologic cross-sectional surveys of a female Lupus cohort, we documented that patients with higher disease activity, and especially those with active renal disease (i.e., Lupus Nephritis, LN) had high-titer serum IgG-responses to 1/8 RG strains reported to have been obtained from healthy donors ^4^. Reactivity with this RG type strain, CC55_001C (which we termed RG2)(Figure 3), was predominantly against a novel lipoglycan (LG), but such antibody responses were infrequent in patients with low disease activity and undetectable in other immune-mediated glomerulopathies or healthy female controls ^4^.

To investigate whether LGs with such antigenic determinants induced LG-specific antibody responses because they are potentially expressed within the gut of Lupus patients, (i.e., expressed by Lupus derived RG strains), we performed immunoblots with the serum of the S47 patient, one of the Lupus strain donors, obtained at the time of clinical flare (Figure 4A-E).

**Figure 4.**
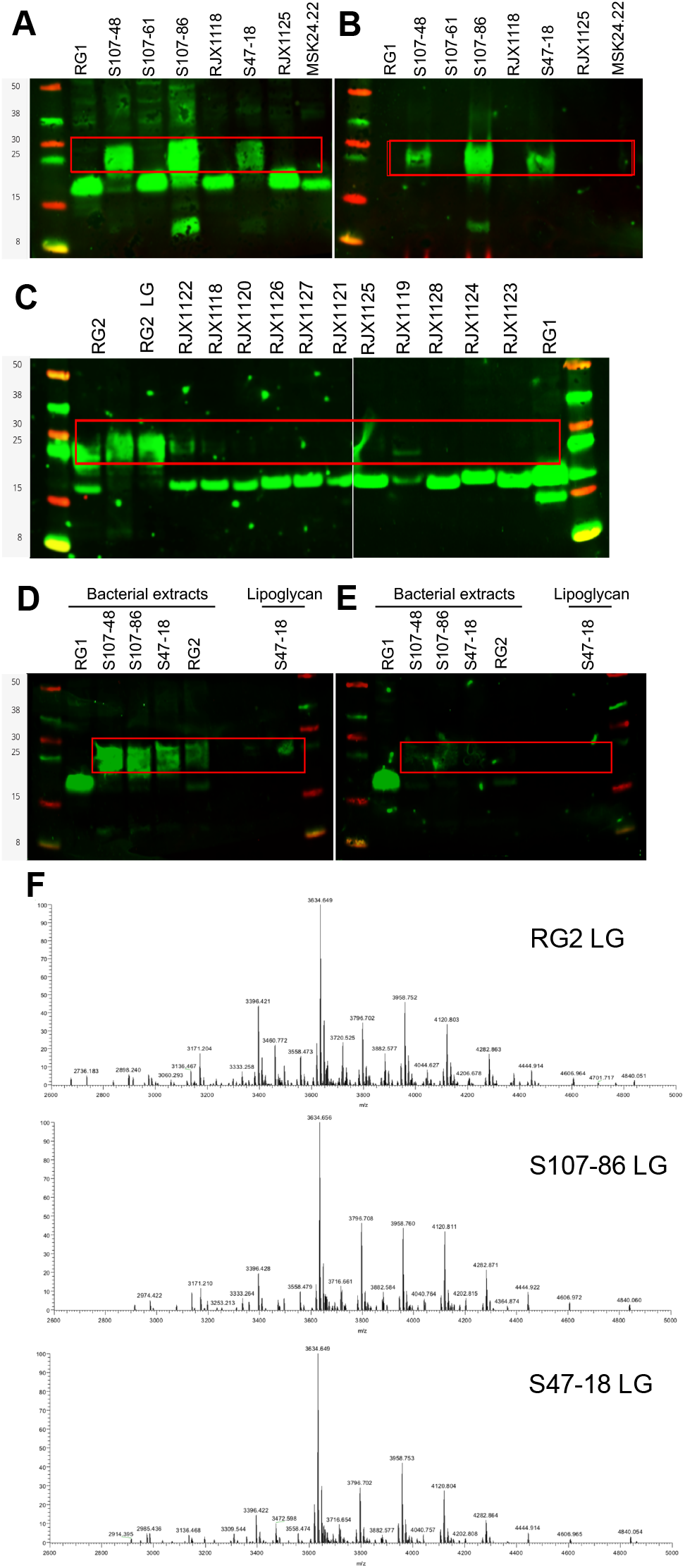
Lipoglycan in different RG strains obtained from Lupus donors during nephritis flares are antigenically and structurally related. A) Molecular components of different RG strains were separated by polyacrylamide gel electrophoresis, transferred to membranes, and immunoblotted with serum IgG from patient S47, obtained during a disease flare. Strains isolated from Lupus patient S107 (S107-48, S107-61 and S107-86), and from patient S47 (S47-18), while RJX1118 was from an antibiotic-treated infant and RJX1125 strain was from a patient with IBD ^14^, and RG1 and MSK22.24 strains were from healthy donors. B) Aliquots of these same bacterial extracts were treated with proteinase K, which reveals the immune recognition by Lupus serum IgG with persistent oligobanding non-protein antigens. C) Nine different RG strains from IBD patients, and two RG strains from antibiotic-treated infants, do <u>not</u> contain the immunoreactive oligobanding lipoglycan found in RG2 extract and purified RG2 lipoglycan. These strains and their genomes described in ^14^. Non-reactive RG1 is also from a healthy adult. D) Immunoblot reactivity of S47 serum IgG-antibodies were reactive with extracts of five RG strains, and the lipoglycan purified from Lupus S47-18 strain. E) A replicate immunoblot was performed after preincubation of the S47 sera with 4ug of purified S47-18 LG, which resulted in complete inhibition of Lupus serum IgG reactivity with oligobands of comparable MW in S107-48, S107-86 and S47-18 extracts and the purified S47-18 LG. The lower MW band in RG1 from a healthy donor, which was previously shown to be protease sensitive, was unaffected. Lipoglycan oligobands migrate to an area delineated by the red boxes. F) Mass spectroscopy-based structural analyses were performed after hydrazine treatment that cleaves off all ester-bound residues, such as fatty acids, as previously described ^4^. Compared with the MS spectrum of native RG2 LG the de-O-acyl RG2 LG displayed significantly decreased complexity in native LG3, the signals at 3380.395 Da, 3394.410 Da and3408.426 Da shifted to the signal of 2931.966 Da in the spectrum of de-O-acyl LG3 (start of series Bde-O-acyl, due to the removal of two fatty acids with combined chain length from C30:0 to C32:0. These changes reflected mass differences of 448.43 Da, 462.44 Da or 476.46 Da, respectively. For this series, molecules with varying number of hexoses were also detectable, as the signal pattern displays mass differences of 162.05 Da. A similar pattern was observed in the lower molecular range between 2000 and 2800 Da for-series Ade-O-acyl, for example, signalsat 2050.663 Da, 2212.716 Da, 2374.768 Da and 2536.820 Da, representing the corresponding de-O-acylated molecules present in series Anative. The core glycan of the purified LG from these three RG strains has a remarkable conservation of overall spectral peaks.

Herein, serum IgG-antibodies were reactive with moieties in extracts of diverse RG strains (Figure 4A), and after protease digestion of these extracts there was persistence of IgG-reactive oligobands of the same 25-30 kilodalton distribution expressed by RG strains, S107-48, S107-86 and S47-18, from the two LN patients. Yet, these antibodies were non-reactive with a RG type strain, ATCC29149 (here termed RG1) or with the MSK24.22 strain ^15^ that were both from healthy donors, and also non-reactive with nine RG strains from IBD patients (RJX1125, RJX1122, RJX1120, RJX1126, RJX1127, RJX1121, RJX1128, RJX1124, RJX1123). Notably, the RJX1120, RJX1121 and RJX1123 strains, which were devoid of detectable lipoglycan, are reported to express a capsular polysaccharide with anti-inflammatory properties ^16^. In addition, two RG strains from antibiotic-treated infants with type 1 diabetes mellitus (RJX1118, RJX1119) all with known genomic sequences (Figure 3B)^14,17^, were also devoid of these IgG-reactive antigenic oligobands (Figure 4A-C). Such antibody reactivity was also absent against the S107-61 strain, but this genome displays modifications attributable to transposon activity (Supplementary Figure 9), which could disrupt genes responsible for LG production.

From evaluations of the relatedness of LG antigens by immunoblot, oligobands were detected in the extracts of the Lupus strains, S107-48, S107-86 and S47-18, that were of the same MW distribution as found in LG purified by hydrophobic interaction column fractionation from the Lupus S47-18 strain (Figure 4A&B). Strikingly, preincubation of this same Lupus sera with LG purified from RG2 strain ^4^ resulted in complete inhibition of reactivity with oligobands, in the RG extracts, and the LG purified from the Lupus RG strain, while the lower MW, protease-sensitive band in the RG1 type strain was unaffected (Figure 4D&E).

To provide a global overview of the distribution of the ion signals as a function of the mass-to-charge ratio, mass spectroscopy (MS) was performed on the LG purified from three independent RG strains (Figure 4F). Strikingly, the purified LG of the RG2 strain and the Lupus strains, S47-18 and S107-86, displayed MS peaks that were (near) identical (Figure 4F). These findings are pathognomonic of the microheterogeneity of bacterial glycans that share dominant structural features, and the differences documented were minor and could have been due to technical variation alone. Taken together, these findings document the conservation of LG, with conserved antigenic determinants, in RG strains isolated from blooms in active LN patients.

### Post-immunization murine antibodies recognize structurally conserved LGs from Lupus RG strains

To provide an independent approach to investigate the antigenic diversity expressed by different RG strains, we generated murine monoclonal antibodies by RG bacterial immunization (see methods). By ELISA, both the mAb 33.2.2 and mAb 34.2.2 strongly react with the purified LGs from the S47-18 strain used for the immunization boost and the purified LG from the RG2 strain (Figure 5A). Both mAbs were reactive with oligobands of the same MW in bacterial extracts from the RG2 strain and Lupus strains, S47-18, S107-48 and S107-86, that were isolated from two different LN patients (Figure 5A). However, these antibodies were non-reactive with the RG1 extract (Figure 5A).

**Figure 5.**
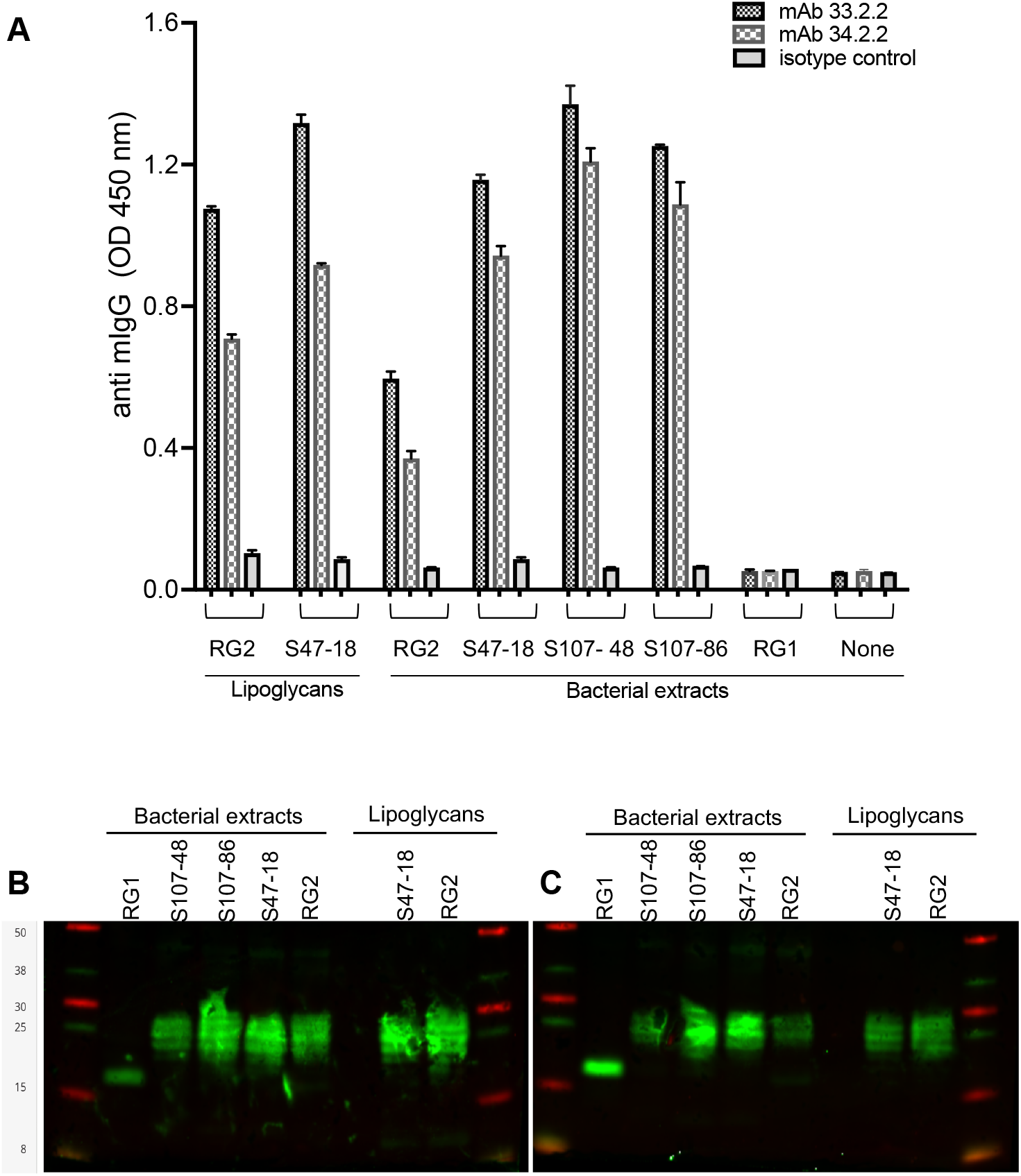
Reactivity of post-immunization murine monoclonal antibodies is restricted to conserved cross-reactive determinants on the oligobands of protease-resistant lipoglycans from RG strains derived from clinically active LN patients. A) Direct binding ELISA demonstrates reactivity of both mAbs, 33.2.3 and 34.2.2, with the purified RG2 and S47-18 LGs, and bacterial extracts from the RG2 strain and Lupus-derived strains; S47-18, S107-48 and S107-86. In this assay, LG or nuclease-treated RG strain extracts are precoated overnight onto the well, then after blocking mAbs, or isotype control at 1 ug/ml are incubated for 4 hrs, then washed and developed (see methods). B) & C) Immunoblots with either of two murine monoclonal antibodies detect antigenically related LG oligobands in extracts of *R. gnavus* strains isolated from active Lupus nephritis strains. In each panel are shown samples of purified lipoglycan from the Lupus S47-18 strain, and the RG2 strain of *R. gnavus*. At left extracts of whole bacteria are shown for the RG1 strain (from a healthy donor), and the Lupus derived strains; S107-48, S107-86, S47-18 and the RG2 strain.

The commonality of these recognized non-protein antigens was confirmed based on reactivity by two mAbs. These also recognized oligoband antigen of the same MW in extracts of the Lupus RG strains from two patients, S107-48, S107-86 and S47-18, as well as the index RG2 strain. These mAbs also bound purified LG from both the S47-18 strain and RG2, while there was no reactivity with the extract of RG1, a type strain from a healthy donor (Figure 5B&C). Notably, the mAb 33.2.2 is of the IgG2a subclass while mAb 34.2.2 is of the IgG1 subclass, which strongly suggests these are products of B-cell clones that express independent antibody gene rearrangements that are convergent in encoding for binding of RG LG antigen-specificity. Taken together, these findings document that RG strains, which colonize different Lupus patients, express structurally related (based on mass spectroscopy) highly immunogenic, cell wall-associated LGs with conserved cross-reactive antigens, and these LGs are recognized by antibodies that spontaneously arise within human Lupus-affected immune systems, and by post-immunization murine monoclonal antibodies. By contrast, these LG do not appear to be expressed by RG strains from healthy donors, and RG strains from IBD patients that in some cases are reported to express forms of capsular polysaccharides ^16,18^.

### High-titer antibody responses arise against structurally conserved RG lipoglycans of Lupus strain associated blooms

To investigate the Lupus host immune response to the expansion of RG strains overtime, we studied the available longitudinal sera from three Lupus patients (S47, Figure 6A-D; S61, Figure 6E-H; and S78, Figure 6I-L), which included serum samples obtained at the time of clinical LN flare (see Figure 2C). Serum IgG-reactivities, at multiple dilutions, were assessed for binding interactions with whole bacterial extract of RG2, the index RG strain first shown to contain an immunogenic LG (Figure 6A, E, I) ^4^, with comparisons of IgG-reactivities with purified LG from this same RG2 strain (i.e., RG2 LG) ^4^ (Figure 6B, F, J). The near identical reactivity patterns of whole bacterial extracts and purified LG, confirm the high immunogenicity of the LG within the RG2 bacterial extract. The same patterns of IgG-reactivities were documented with the structurally-related LG from the S47-18 strain (S47-18 LG) obtained from the S47 LN donor (Figure 4D&E, 5A&B), although for this LG there were uniformly stronger binding interactions with the IgG in sera from each of the three LN patients (as IgG-binding curves for S47 LG were shifted up and to the right, Figure. 6C, G, K). By contrast, there was little or no detectable reactivity with the unrelated LPS glycan from a *Pseudomonas* species (Figure 6D, H, L) that is not known to bloom within these subjects. Taken together, these data reveal high-titer Lupus host LG-specific serum antibody reactivities, with peak levels in Lupus patients S47 and S61 at the time of disease flare, which was also concordant with an RG bloom in abundance. For patient S78, the antibody titers were relatively invariant, and substantial disease activity was documented at most visits (Figure 6I, J, K). Notably, the strongest antibody reactivity of the S47 serum antibody reactivity was with the purified LG from an RG strain isolated from a S47 fecal sample (Figure 6C), which was obtained at time of disease flare and RG bloom. Importantly, parallel highest IgG-reactivity was also found in the sera of the other two Lupus patients (Figure 6G&K). These findings document the intertwined nature of Lupus disease flares, which involve immune-complex mediated pathogenetic pathways ^19^, and at time with host immune responses to a novel RG Lupus strain-associated LG (Figure 6).

**Figure 6.**
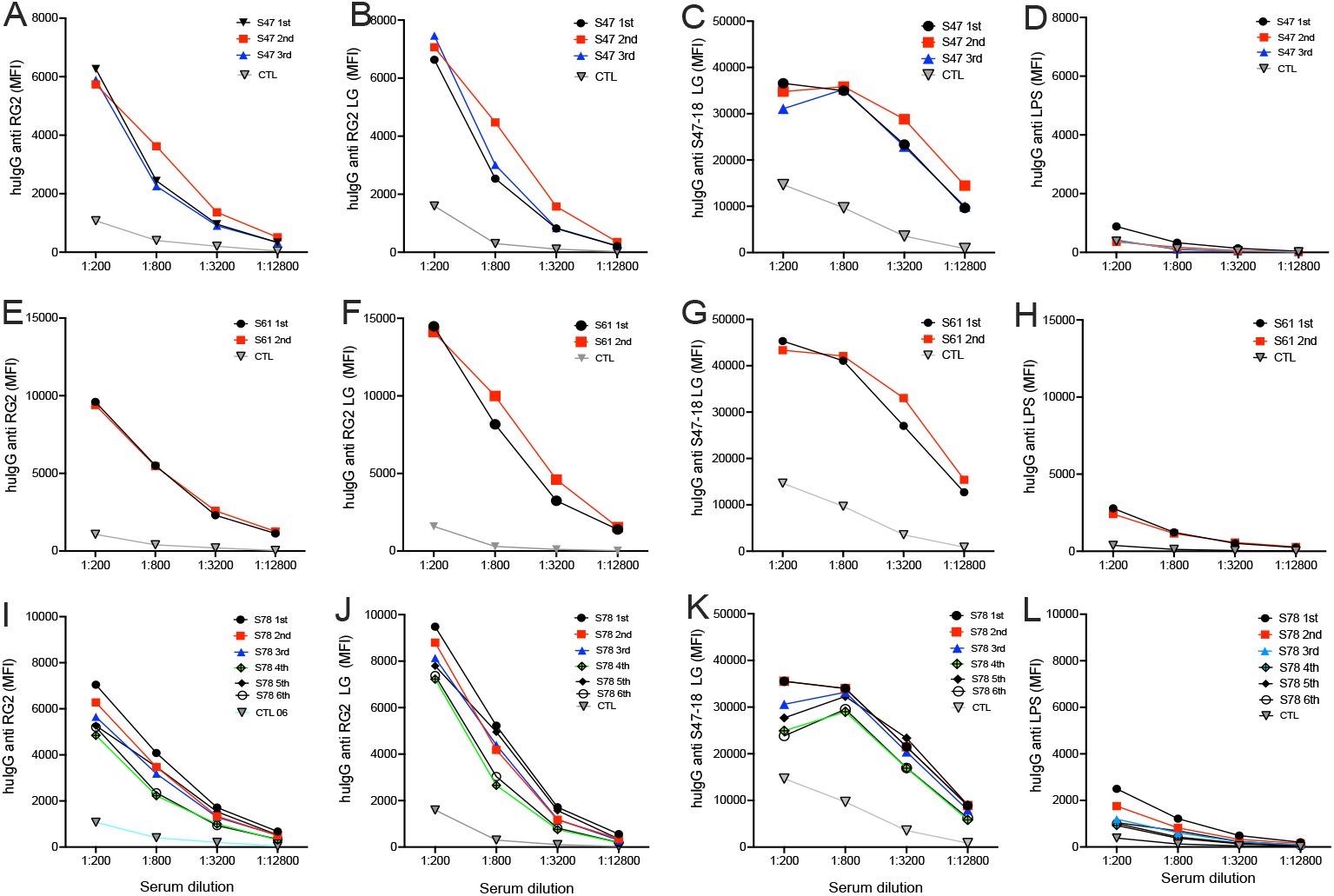
Serum IgG antibodies to Lupus RG strain LGs, parallel gut community abundance, peak with RG blooms and concordant disease flares. To investigate the Lupus host immune response overtime to colonization with RG strains, we studied IgG anti-RG responses longitudinally obtained sera from three Lupus patients. A-D) Results for IgG responses for patient S47. E-H) For patient 61. I-L) For patient S78. A, E, I) Serum IgG-reactivity at multiple dilutions was assessed for binding to whole bacterial extract of RG2, the index RG strains that contained the first identified immunogenic LG. B, F, J) Comparisons with purified LG from this same RG2 strain (i.e., RG2 LG), which show very similar reactivity patterns, confirming the high immunogenicity of LG within the whole bacterial extract. C, G, K) Reactivity with the structurally related LG (see Figure 4C) from the S47-18 strain (S47-18 LG) from the S47 LN donor, the same relative reactivity patterns are seen, although uniformly stronger levels) were seen with sera from each of the three patients (i.e., IgG-binding curves for anti-S47-18 LG are shifted up and to the right). D, H, L) By contrast, there is little or no reactivity with the LPS glycan from a *Pseudomonas* species, which is not known to be a blooming bacterial species within these subjects. In each panel, reactivity with serum from a representative healthy female control is shown (CTL). Samples obtained sequentially overtime are indicated accordingly. Studies performed with custom bead based MBI array (Luminex), with previously described methods ^4^.

## Discussion

Herein, we report longitudinal studies demonstrating temporal instability within Lupus gut microbiota communities from which blooms of RG arise concordant with Lupus disease flares (Figures 1&2). Notably, the level of variance overtime within these communities did not correlate with; numbers of samples from a donor, the timespan of sampling, or the features of the disease activity. This variance was neither correlated with maximum SLEDAI disease score during the period of study nor span from lowest to highest SLEDAI score. Furthermore, the greater variance in Lupus patients also did not appear to correlate with medication taken (Supplementary Tables 1 and 3). Hence, the inherent instability of the gut microbiota in Lupus patients appears to be a feature inherent to the disease.

Whereas RG was consistently documented only at low stable levels overtime in healthy subjects that was not the case in SLE. By contrast, in a subset of the Lupus patients (5/16), representing 4/9 LN patients, the dynamic RG blooms were strikingly concordant with periods of greatly increased disease activity, which in almost every case was manifest as flares involving renal disease. However, one patient (S61) had no history of renal involvement, her RG gut bloom was concurrent with an arthritis flare involving multiple joints. These RG bloom-associated disease flares occurred in White, African-Americans, Asian and Hispanic patients (Supplementary Table 1)^4^, which may suggest that, while RG blooms trigger flares of Lupus disease activity, the patterns of organ involvement reflect patient-specific factors—there are likely genetic factors that predispose to renal involvement, akin to differences in murine Lupus strains that are susceptible to, or are protected from, nephritis. Autoantibody profiles may also be contributory, as certain types of IgM-autoantibodies may be protective from renal injury ^20^, although antibody levels to RG were not here evaluated.

The RG strains recovered from Lupus patients with blooms produce a novel lipoglycan, with pro-inflammatory properties, and structural features that are conserved based on mass spectroscopy analysis, and conserved antigenic determinants recognized by murine monoclonal antibodies. The LG purified from the index RG2 strain has been shown to have N-acetyl galactose (GalNAc) residues and mannose capping as well as a diacetylglycerol core ^4^ akin to the LG of *Corneybacterium glutamicum* ^4,21^. Foremost, these RG lipoglycans are also highly immunogenic, inducing high-titer systemic IgG responses, which parallel RG abundance in cross-sectional and longitudinal studies, especially in LN with active disease (Figure 6). Notably, in cohorts across the US ^4^, and high-titer IgG antibodies to RG LG were also documented in active LN in a European cohort ^22^, suggesting this linkage of SLE pathogenesis and hot colonization with lipoglycan-expressing RG strains has no simple geographic restriction.

Emerging evidence further suggest that subclinical disturbances in gut-barrier function, found in other conditions ^23^, may be common in Lupus patients ^4,24^. Indeed, in a murine Lupus model, a gut pathobiont was shown to translocate from the intestine and cause an inflammatory condition ^25^. Furthermore, fecal transplants from mice with Lupus-like glomerulonephritis induce an inflammatory and autoimmune state in otherwise non-autoimmune germ-free C57BL/6 mice ^26^, which clearly demonstrated the pathogenic potential of some gut microbiota communities. Highly relevant, murine colonization with RG strains that expressing this lipoglycan, but not the control RG1 strain, induced severe gut leakiness, and production of antibodies to the lipoglycan, and to native DNA^27^, a hallmark of Lupus. Within the aforementioned cross-sectional studies, levels of anti-RG2 antibodies also significantly correlated with serum levels of IL-6 and type I interferon, TNF-α, IL-10 ^4^. Taken together, we surmise that anti-RG responses arise in concert with autoantibody production, which may be a consequence of bacterial factors contributing to the stimulation of autoreactive B cells and disease-specific autoantibody production in predisposed hosts. Indeed,

In the serial blood samples from Lupus patients, antibody responses to RG, and specifically for the RG lipoglycan, paralleled the level of RG gut abundance (Supplementary Figure 5). In our earlier report, levels of antibodies to RG2 strain (CC55_001C) that is dominated by recognition of LG antigen, inversely correlated with C3 and C4 levels, which was suggestive of an associated immune complex disease ^4^. In addition, anti-RG2 levels correlated with IgG anti-dsDNA levels. Nonetheless, we are careful to not overinterpret the results. Firstly, while RG abundance was measured in fecal samples, the relationship is currently unknown with RG abundance at other intestinal sites, such as in ileum and along mucosal surfaces implicated in luminal antigen transport in the “leaky gut”, for translocation of other bacterial factors into the systemic immune system and for modulation of host immune responses.

RG is a member of the Lachnospiraceae family that plays pleotropic roles in host metabolism and immunity (reviewed in ^28^), including for bile acid metabolism ^29^ and the production of the short-chain fatty acids (SCFAs) that aid immune regulation ^30^, has otherwise generally been associated with pro-homeostatic communities ^16^. It is therefore important to note that based on whole genome differences, as well as 16S rRNA sequence variations, RG is genetically distinct from all other members of this family, and from other Blautia isolates ^15^. Importantly, the RG strains in the blooms in Lupus appear to be different than the RG expansions that have also been documented in independent cohorts with IBD ^14,31^, including patients with IBD associated spondyloarthritis in whom RG abundance correlated with joint disease activity ^31^. Yet, IBD is a disease of the bowel in which RG expansions may be directly involved in tissue injury inherent to the disease, and examination of 9 RG strains from IBD patients found no detectable lipoglycan (Figure 4A-D). IBD patients do not have prominent associations with autoantibody production or immune complex disease, while susceptibility based on Class I HLA B27 alleles has pointed towards very different mechanistic roles of the gut microbiome with IL-17/23 associated pathology ^32^. By contrast, while systemic autoantibody production and immune complex disease are integral to the pathogenesis of Lupus, clinically apparent colitis is not a common complication ^33^.

The limitations of the current clinical single-center observational studies are that samples were collected during routine clinical care and not at strict intervals. We, therefore, could have underestimated the true occurrence of disease flares with temporally linked RG blooms. As we were concerned that medications could have effects on the metabolism and proliferation of commensal species, whenever possible we therefore obtained fecal samples before modification of treatment regimen. Yet, human-targeted medications are not known to have effects limited by bacterial family, genus or species ^34^ and the most commonly used therapeutic agents, corticosteroids and hydroxychloroquine, are reported to have little or no effect on the human gut microbiome ^35^. The design of the current studies was also inadequate to disentangle the potential influence of diet or unreported OTC medications from those integral to Lupus itself. In addition, we could not consider whether gut microbiota instability predates clinical disease. While key structural and antigenic features of RG lipoglycan have now been documented to have remarkable conservation between different LN-derived RG strains (Figure 4), much remains to be defined in the structural and compositional nature of RG LGs. The responsible genes and molecular enzymatic pathways responsible for assembly also need to be identified.

Our findings support the notion that RG expansions can directly contribute to Lupus disease flares. This does not exclude the potential involvement of other pathobionts, or contractions of “protective” species such as *Faecalibacterium prausnitizii* that also occur during active Lupus disease ^4^. Overall, our findings appear to reflect the same altered gut community patterns documented to occur in Chimpanzees after Simian Immunodeficiency Virus (SIV) infection, in which gut microbiota community instability arises and transient spikes/blooms newly emerge ^36^. This gut community (dys-) regulation is likely influenced by CD4+ T cells that are selectively deleted by SIV ^37^, and there may be parallels in SLE patients who are known to have generalized defects in T-cell regulation ^38^. Moreover, while in health CD4+ T cells play central roles in the local maintenance of the gut barrier ^39^, the implications of human Lupus-associated T-cell dysregulation on gut bacterial expansions and altered intestinal permeability have not been examined.

In conclusion, we must now consider whether in a susceptible Lupus patient, RG blooms can stoke systemic inflammation and tissue injury by a post-intestinal bloom autoimmunity syndrome. Emerging evidence suggest that the pathogenic potential of an RG strain is influenced by the production of specific classes of bacterial glycans, such as the lipoglycan expressed by Lupus strains ^4^. By contrast, many RG strains are reported to have genes that encode for glucose-rhamnose capsular polysaccharide that stimulates innate immune cells to produce inflammatory cytokines (Henke *et al*. 2019), while a strain from a healthy donor (i.e., RG1) as well as IBD-derived RJX1120, RJX1121, RJX1123 produced a “tolerogenic” capsular polysaccharide that may provide homeostatic benefits to the host ^16^. However, our immunochemical surveys found these other strains were devoid of lipoglycans related to the product of the Lupus strains (Figures 3&4). As multiple RG strains have been isolated from a single donor ^15^, disease flares could therefore be in part a consequence of ecological shifts in the representation within a community of individual RG strains that differ based on their associated sets of genes, and the specific cell wall-associated glycopolymer products, that determine their pathogenic potential.

The next steps toward understanding how RG blooms affect Lupus pathogenesis will require the in-depth characterization of RG colonies directly isolated from Lupus patients in different cohorts, as we strive to understand whether, in a susceptible host, certain RG strains are inherently nephritogenic, and we will seek to identify the genes in the RG genome that encode for the responsible virulence factors.

## Materials and methods

Studies were performed under the supervision of the NYU Institutional Human Subjects Committee, and all subjects provided written informed consent. Exclusion and inclusion criteria for patients and healthy controls have previously been described ^4^. These specific criteria, other aspects of the clinical studies, and the collection and characterization of the gut microbiota is described in the supplement (Table 1).

### Isolation, characterization, genome sequence determination of RG colonies

To identify and recover *R. gnavus* colonies, fecal samples were streaked onto TSBA or BHI plates and grown under anaerobic conditions. Individual colonies identified based on morphology and growth characteristics of the strains ATCC29149 (termed RG1) and CC55_001C (termed RG2), then sub-streaked and sub=cultured. From each colony genomic DNA was recovered with the power soil (Qiagen™) kit, according to the manufacturer’s instructions, quantified on a Nanodrop 1000 (Thermo Scientific). To assess total 16S rRNA gene, PCR assays were performed with the T100 thermocycler (Bio-Rad™) using the primers:

For all bacteria ^40^:

UniF340(5’-ACTCCTACGGGAGGCAGCAGT-3’)

UniR514(5’-ATTACCGCGGCTGCTGGC-3’). 94°C for 3 minutes, followed by 35 cycles of 94 °C for 45 seconds, 50°C for 1 min; and followed by extension at 72°C for 10 minutes and 4°C hold.

The RG species-specific 16S rRNA was determined with the previously reported oligonucleotide primers ^40^:

Fwd 5’-GGACTGCATTTGGAACTGTCAG-3’

Rev 5’-AACGTCAGTCATCGTCCAGAAAG-3’

for the following cycles: an initial 94°C for 3 minutes, followed by 35 cycles of 94 °C for 45 seconds, 58 °C for 1 min; and followed by extension at 72°C for 10 minutes and 4°C hold.

Colonies of interest were named based on the Lupus patients that provided the faecal sample of origin, S47- and S107-, which were named according to the donor source. DNA from each isolate was then subjected to whole genome sequencing, using both NextSeq 550 Illumina and PacBio technologies. Alignments with the genomes of RG1 and RG2 strains documented that the assignment to the RG species with genomes that affirmed that independent non-identical RG strains (see Supplementary Figure 9). Raw sequence data and assemblies have been deposited in NCBI under BioProject PRJNA82122). Purifications of the bound glycopolymers in the S107-86 and S47-18 RG strains were performed as previously described, from which earlier analysis of the RG2 purified moiety was shown to contain a diacylglycerol- (DAG-) containing lipid anchor characteristic of a lipoglycan (LG).

### Bacterial whole-genome sequence analysis

Raw short-read sequence reads were preprocessed using fastp (Chen *et al*. 2018) version 0.22 using default settings. Preprocessed reads were screened for within-species contamination using ConFindr (Low *et al*. 2019) as well as cross-species contamination using MetaPhlAn (Beghini *et al*. 2021). Preprocessed short reads were assembled using Unicycler (Wick *et al*. 2017) using its conservative mode.

Processing and assembly of PacBio long reads was performed using the PacBio SMRT Link software suite. Assemblies were further assessed by BLAST search (Altschul *et al*. 1990) against the NCBI nt database. Contigs matching *Cutibacterium acnes*, a common skin bacterium, were removed from long-read assemblies, and it was verified that the remaining contigs mapped to corresponding uncontaminated short-read assemblies Genome assemblies were annotated using Prokka (Seemann 2014). The BEDTools software suite (Quinlan and Hall 2010) was used to divide genome assemblies into 1kbp windows and to analyze GC content per window. Each window was compared to RG1 (NCBI accession GCF_009831375.1) and RG2 (GCF_000507805.1) using BLAST (Altschul *et al*. 1990), keeping only alignments with E values below 10^−20^. Visualizations of each genome assembly were produced using the circlize R package (Gu *et al*. 2014).

All genome assemblies linked to the species [Ruminococcus] gnavus (taxonomy ID 33038) (based on an older nomenclature) in NCBI RefSeq (accessed March 10, 2022) were downloaded for comparison to the newly generated assemblies. One of these (accession number GCF_020538135.1) was found to be highly divergent from the others (with average nucleotide identity < 90%) and was excluded from further comparisons. Assessment of gene content across assemblies was performed using Orthofinder (Emms and Kelly 2019). The resulting presence/absence matrix of orthogroups was used to generate pairwise Jaccard dissimilarities between isolates using the vegan R package (Oksanen *et al*. 2020). Snippy (Seemann 2015) was used to produce a core genome alignment, using the downloaded RefSeq genome assembly for strain RG1 as reference (NCBI accession number GCF_009831375.1). A phylogenetic tree was inferred from the core alignment using RAxML (Stamatakis 2014) with a GTRGAMMA model and 100 bootstrap replicates.

### Lipoglycan purification and reporter cell assay

RG strains were individually expanded in an anerobic biofermeter in chopped meat media to stationary phase and 300 ml were pelleted, and used in an established extraction procedure designed to isolate lipoteichoic acids from Gram-positive bacteria ^41^. Briefly, to purify the cell wall lipoconjugates, bacterial cells were disrupted with a French press, then the precipitate was removed by ultracentrifugation. The supernatant was subjected to butanol-water extraction and the water-soluble component then passaged over a Hydrophobic Interaction Chromatography (HIC) to isolate lipoglycan-containing fractions. Following our previously described procedure ^42^, hydrazine treatment was performed to yield de-*O*-acylated lipoglycans. These substances were characterized by MS as earlier described ^4,42^

### Immunoblotting

Electrophoretic separation used Bis-Tris mini gels (Novex, Thermo Fisher) with bacterial extracts loaded at the same concentration, then transferred to membranes, which were incubated with sera diluted at 1:100, and incubated overnight at 4°C. For detection, anti-human IgG biotin conjugated (Jackson ImmunoResearch Labs, USA) was added and developed by IRDye^®^ 800CW Streptavidin (LI-COR^®^).

### Generation of LG-specific murine monoclonal antibodies

A commercial vendor (Envigo Bioproducts Inc., Indianapolis) immunized 10 BALB/c mice with extract of the RG2 strain emulsified in complete Freund’s adjuvant and later boosted with lipoglycan purified from the Lupus S47-18 strain, which was emulsified in incomplete Freund’s adjuvant. All LG were purified from an RG strain by a method that included fractionation by hydrophobic interaction chromatography, as previously described ^4^. The spleen from the mouse with the strongest post-immunization was fused with Ig-deficient NS-1 myeloma cells. The spent supernatants subclones were evaluated for IgG-reactivity, which demonstrated highly correlated reactivity with whole extracts of the immunizing RG strain and purified RG lipoglycan, with the subcloned hybridoma cell lines, termed mAb 33.2.2 and mAb 34.2.2.

### Direct binding ELISA

To detect the reactivity of the murine monoclonal antibodies 33.2.2 and mAb 34.2.2 with the different RG strains, the ELISA plates were coated with the bacterial extracts from RG2, S47-18, S107-48, S107-86, RG1 as well as with the purified lipoglycan from the strains RG2 and S47-18. Next, the murine monoclonal antibodies were added at 2 concentrations (at 100ng/ml and 25ng/ml) in duplicate, incubated for 2hrs at RT. Binding was detected with goat anti mouse IgG HRP conjugated at 1:10,000 (Jackson ImmunoResearch), then TMB substrate was added to develop the plate.

### Multiplex Bead based immunoassay

The assays performed as previously described in ^4^. Briefly, serum samples from patients and healthy controls underwent to 4-fold serial dilutions starting at 1:200 to 1:12,800 against a panel of antigens including the bacterial extracts from Lupus strains RG2, S47-18, RG1 as well as the purified lipoglycan from the strains RG2 and S47-18, then detecting using goat anti-human, PE, (eBioscience).

### Statistical analysis

Data are presented as mean ± SD. The Student unpaired *t* test with Welch correction was used in 2-group comparisons of normally distributed data, whereas the Mann-Whitney nonparametric test was used when the normality assumption was not met. Fisher’s exact test was performed to evaluate bivariate associations between categorical variables, or as described. To test for correlations between two variables Spearman test was used. *p-*values were considered significant at <0.05 for two-tailed tests. Prism software Version 9 (GraphPad) was used for all analyses.

## Acknowledgments

We appreciate the assistance of the NYU Immune Monitoring Core supported by NYU-HHC CTSI Grant UL1 TR000038, the NYU the Laura and Isaac Perlmutter Cancer Center support grant, P30CA016087 from the National Center for Advancing Translational Sciences (NCATS), the NYU Genome Technology Core including for *PacBio sequencing that was supported by the NIH Shared Instrumentation Grant 1S10OD023423-01*, and the Microbial Computational Genomic Core Lab. We thank Jeffrey Weiser, Matt Henke and Dan Littman for advice, Ramnik Xavier and Hera Vlamakis, Eric Pamer, and Emma Allen-Vercoe for providing RG strains, and Mimi Kim for biostatistical support. We acknowledge the staff of Envigo Biologics Inc. for production of monoclonal cell lines.

Funding This work was supported in part by National Institutes of Health Grants; R01-AR42455 (GJS), P50 AR070591 (GJS, JPB, AVA), NIAID contract for B Cell Epitope Discovery and Mechanisms of Antibody Protection HHSN272201400019C (GJS), the Lupus Research Alliance, and the Judith and Stewart Colton Autoimmunity Center (GJS). 16S rRNA amplicon sequence determinations and analysis were supported by the P. Robert Majumder Charitable Trust (GJS).

