## Supplementary Figures and Tables for "Lupus disease flares are concordant with immune responses to blooms of lipoglycan-expressing *Ruminococcus blautia gnavus* strains arising from unstable gut microbiota communities"

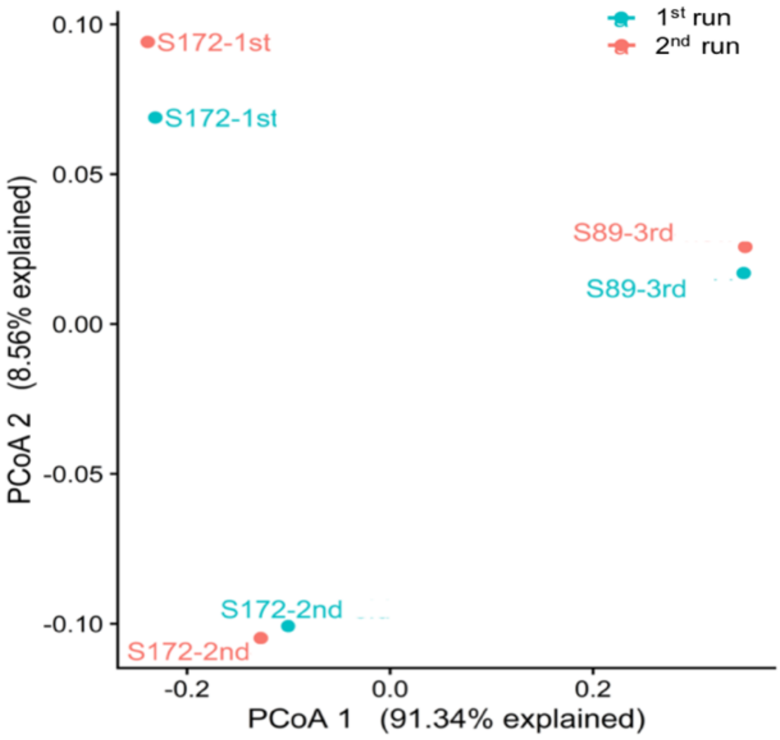


**Supplementary Figure 1. Consistent libraries composition between the two 16S rRNA gene amplimer Miseq batches**. Principal Coordinates Analysis (PCoA) on libraries generated from samples of same individuals/samples sequenced in the two different 16S rRNA sequencing runs ($1^{st}$ and $2^{nd}$) as an inter-run technical control, shows no consistent bias or batch effect.

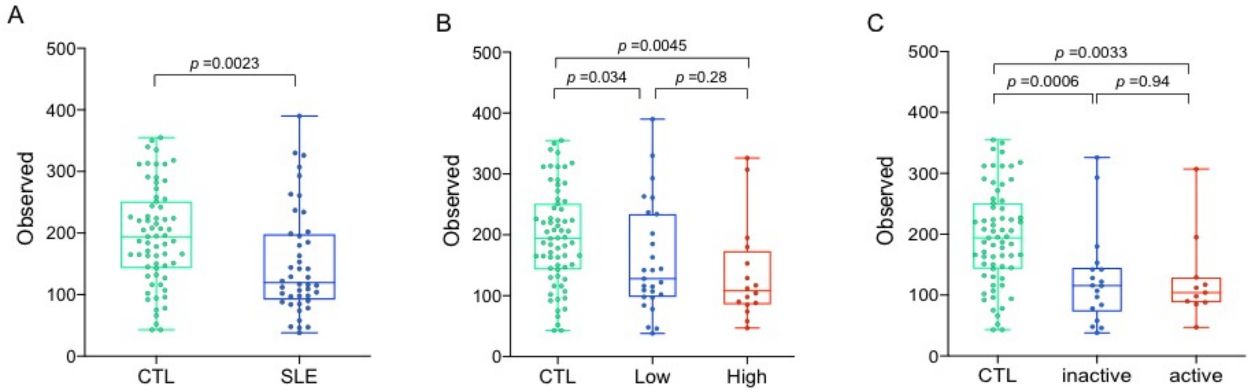


**Supplementary Figure 2.** Dysbiosis in **in SLE microbiota communities**. A) The number of distinct taxa were estimated based on observed ASV, and alpha diversity richness was reduced in samples from Lupus patients (SLE) compared to healthy controls (CTL) (Wilcoxon, $p$=0.0023). B) Compared to CTL, alpha diversity was reduced in Lupus patients with low disease activity (based on SLEDAI score), with even greater contractions in the group with high disease activity, (Wilcoxon, $p$=0.034, $p$=0.0045, respectively). C) Compared to CTL, alpha diversity was reduced in those with inactive renal disease, and further contracted in those with active renal disease (Wilcoxon, $p$=0.0006 and $p$=0.0033, respectively). The purpose was to assess to consider correlations that in an individual subject, might be constant, or vary overtime, these analyses did not consider statistical adjustments for multiple samples from the same patient.

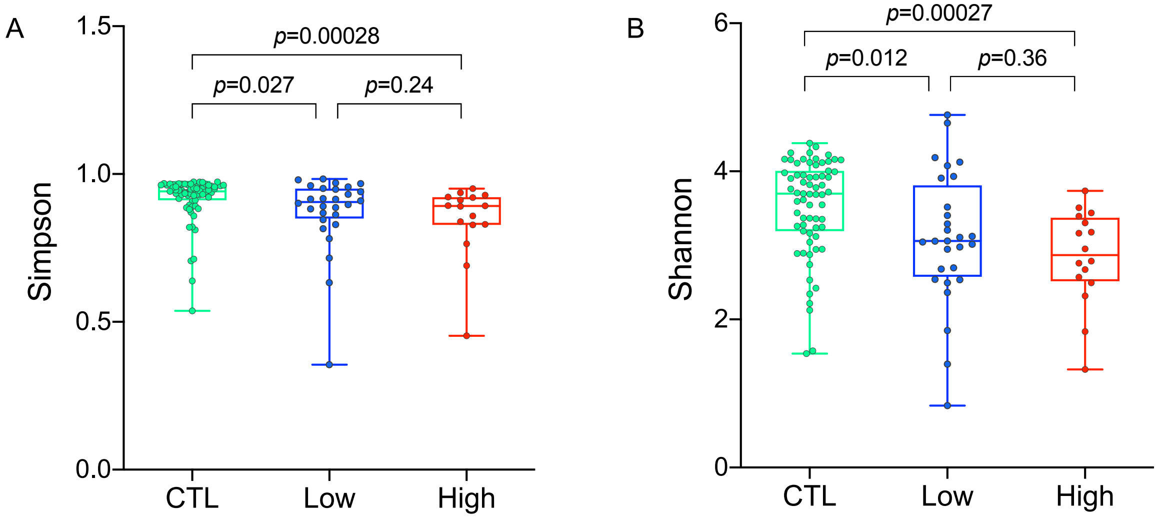


**Supplementary Figure 3. Alpha diversity is reduced in libraries from patients with high Lupus disease activity.** Analyses are as shown in Figure 1. High disease activity is defined as a composite SLEDAI score of $\geq$8.

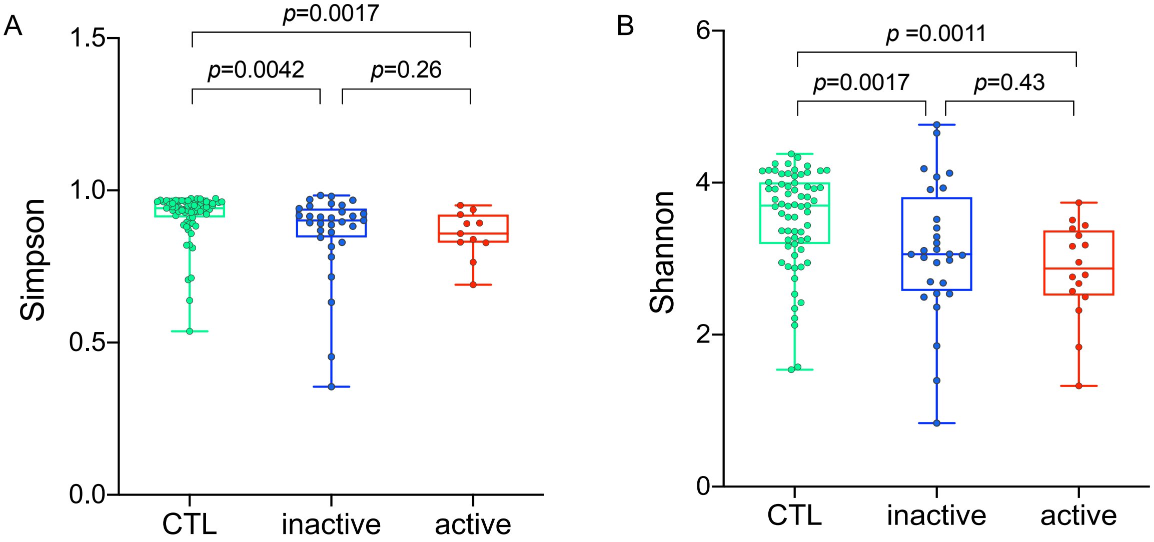


**Supplementary Figure 4. Alpha diversity is reduced in libraries from patients with active renal disease compared to inactive renal disease.** Analyses performed as shown in Fig. 2. Active renal disease was defined by standard clinical laboratory criteria.

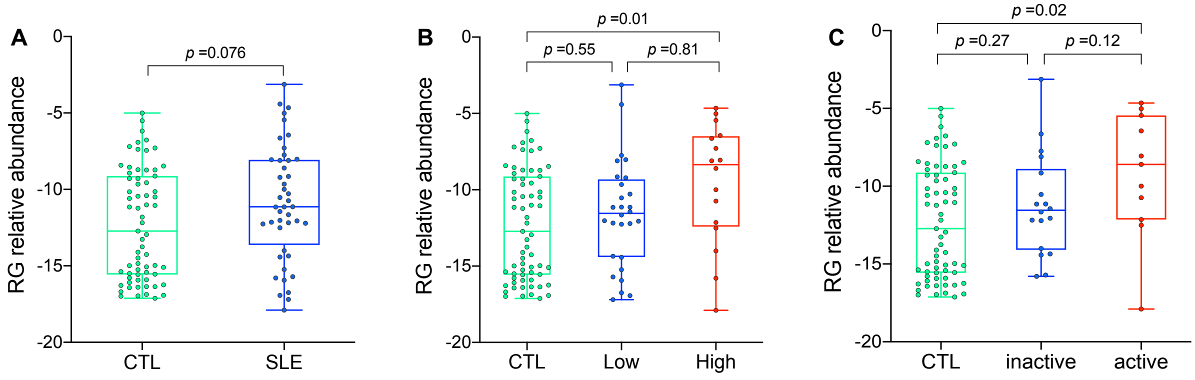


**Supplementary Figure 5. RG expansions occur at the time of high Lupus disease activity and active LN**. A) All samples from SLE patients have a numerical but non-significant trend toward increased RG abundance compared to healthy CTL (Wilcoxon, *p*=0.0760). B) Samples from patients with high disease activity (based on SLEDAI) showed a greater RG abundance, compared to low disease activity and CTL (Wilcoxon, *p* =0.81, *p* =0.01, respectively). C) RG expansions were common in the active LN group compared to healthy CTL (Wilcoxon *p*=0.02), but not significantly different in the inactive LN group Wilcoxon, *p*=0.27, NS. RG relative abundance is shown in log 2 values.

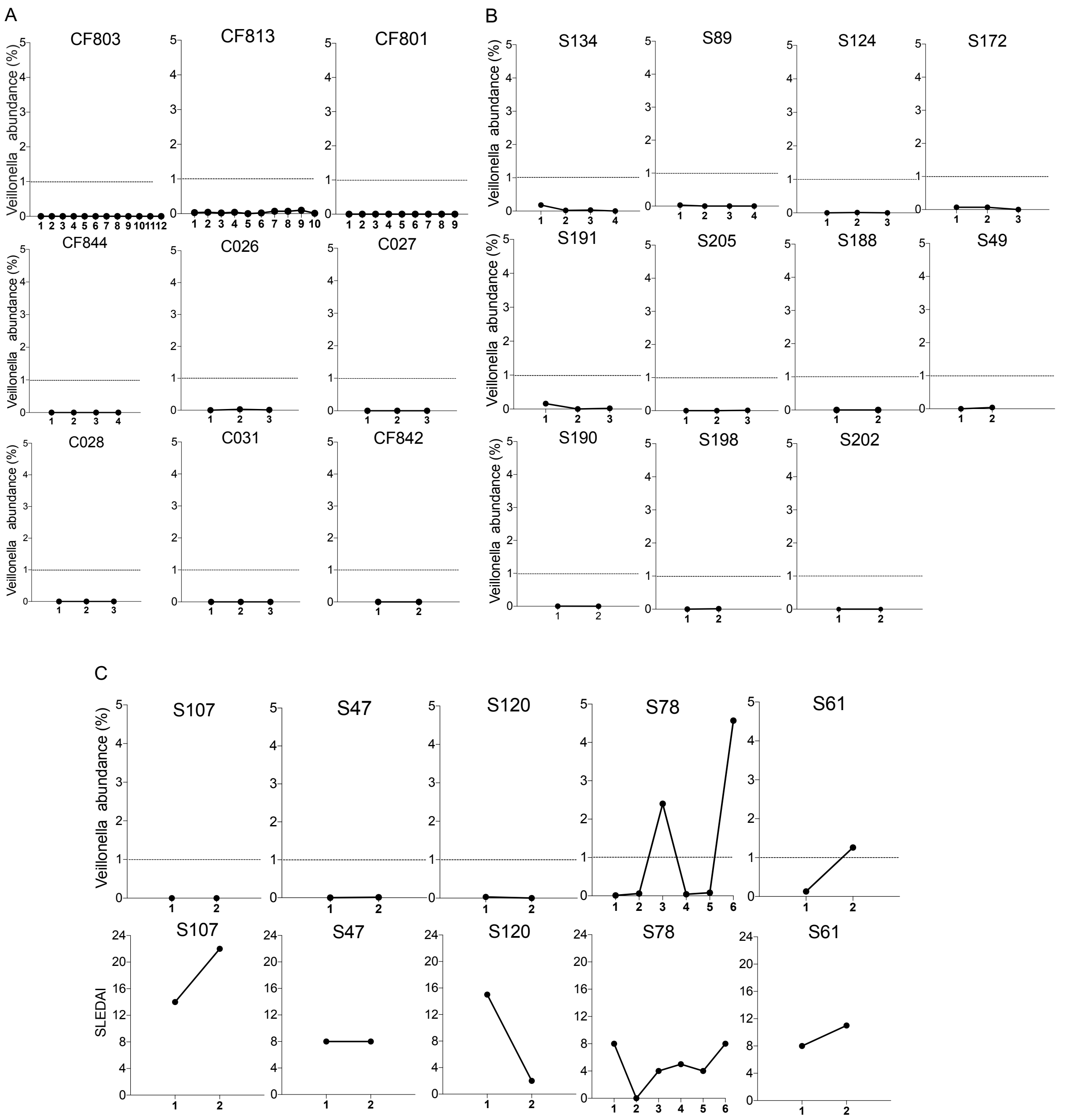


**Supplementary Figure 6**. **Blooms of *Veillonella* do not occur concurrent with Lupus disease activity**. A) *Veillonella* abundance in healthy individuals. B) *Veillonella* abundance in SLE patients. C) *Veillonella* abundance in SLE patients with above-described RG blooms concordant with Lupus disease activity flares. Abundance based on ASV representation in total amplicon libraries.

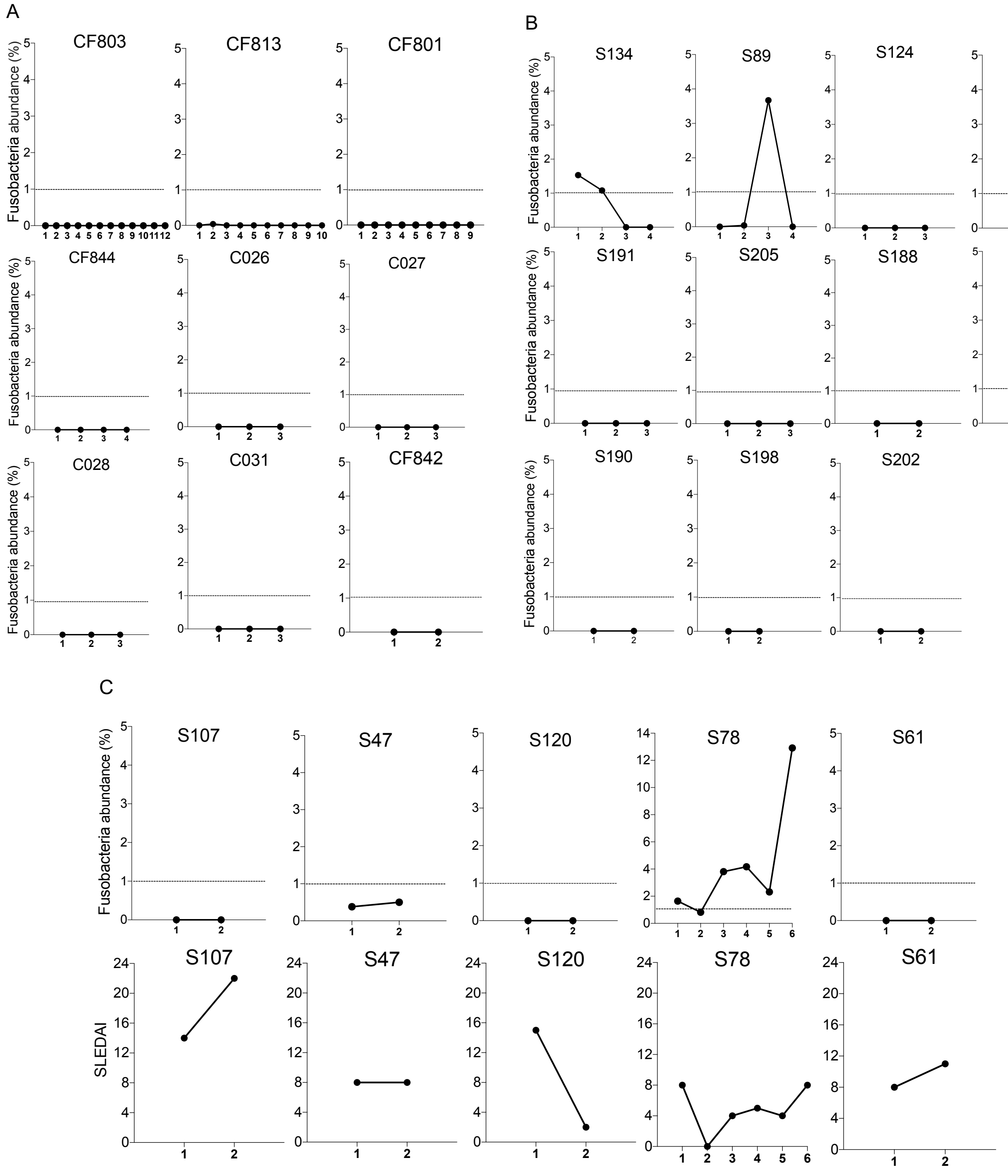


**Supplementary Figure 7**. **Blooms of *Fusobacterium* do not occur concurrent with peak flares of Lupus disease activity**. A) *Fusobacterium* abundance in healthy individuals. B) *Fusobacterium* abundance in SLE patients. C) *Fusobacterium* abundance in SLE patients with above-described RG blooms concordant with Lupus disease activity flares. Abundance based on ASV representation in total amplicon libraries.

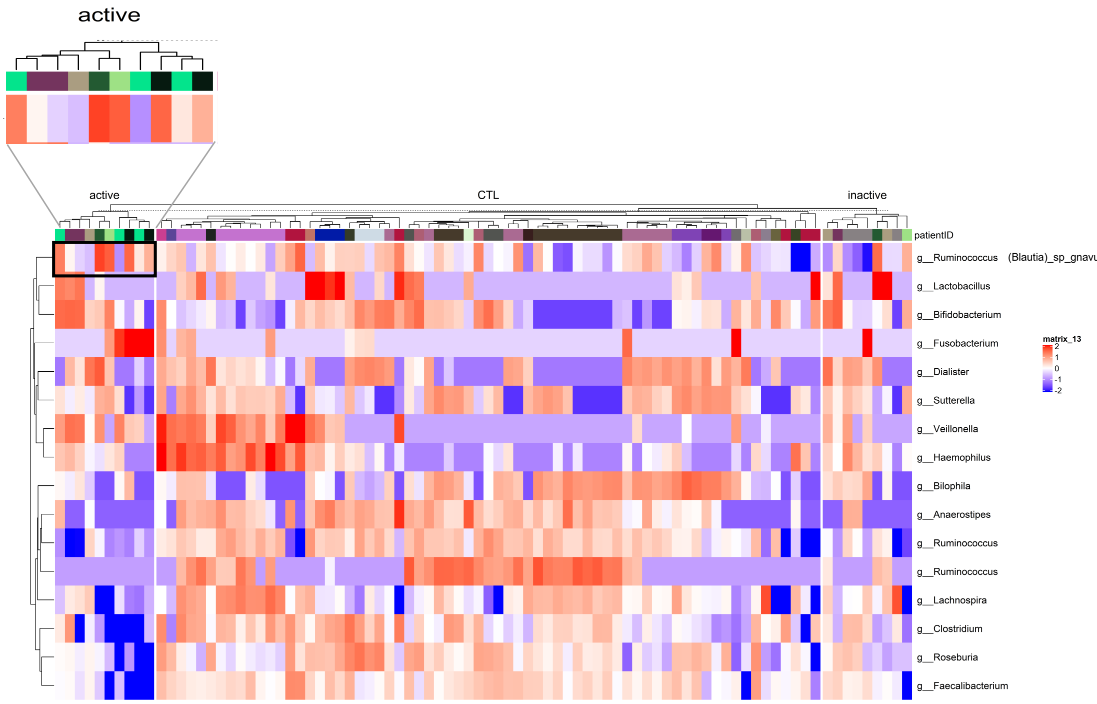


**Supplementary Figure 8. Heat map at the species level shows dominance of *Ruminococcus (blautia) gnavus* in many patients with active LN.** The abundance of the species in the different groups (healthy controls CTL), patients with active nephritis and with inactive nephritis, by using the Z-score. Cut-offs were used that were calculated based on the Z-score. Taxa with mean of relative abundance of greater than 0.001 are depicted, and taxa with DEseq2 padj > 0.1 across all samples. The paucity of *Faecalibacterium* is evidence in Lupus patients.

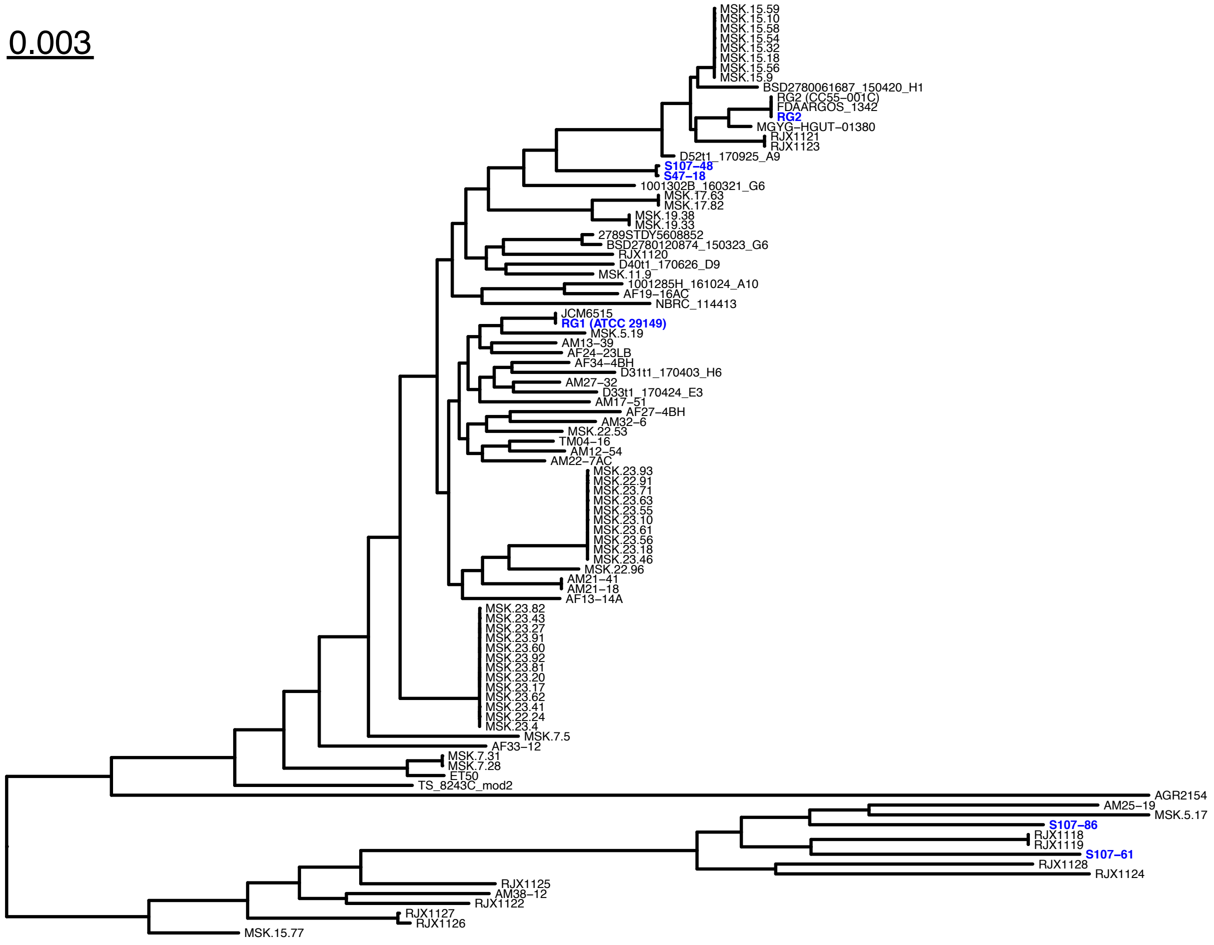


**Supplementary Figure 9.** Phylogenetic tree based on core alignment of *Blautia (Ruminococcus) gnavus* (RG) genome assemblies downloaded from NCBI RefSeq, together with five newly generated genome assemblies. Newly sequenced isolates are shown in blue, along with downloaded RG1 (ATCC 29149) strain for reference.

**Supplementary Table 1A. Demographic, clinical and treatment features of Lupus patients with renal involvement evaluated overtime.**

| Patient ID | Age | Ethnicity | Sample | Collection week span | SLEDAI | Renal ACR | Renal active | MEDs | | R. *gnavus* % |
|---|---|---|---|---|---|---|---|---|---|---|
| | | | | | | | | Prednisone | MMF | |
| S47 | 39 | Asian | 1st | 0 | 8 | 1 | 1 | 0 | 1 | 1.1 |
| | | | 2nd | 256 | 8 | | 1 | 0 | 1 | 2.3 |
| S78 | 38 | White Hispanic | 1st | 0 | 8 | 1 | 0 | 10 | 1 | 0.4 |
| | | | 2nd | 176 | 6 | | 0 | 7.5 | 0 | 0.0 |
| | | | 3rd | 205 | 4 | | 0 | 5 | 0 | 0.2 |
| | | | 4th | 246 | 5 | | 0 | 5 | 0 | 0.0 |
| | | | 5th | 268 | 4 | | 0 | 5 | 0 | 0.1 |
| | | | 6th | 291 | 8 | | 1 | 5 | 0 | 9.5 |
| S89 | 37 | Asian | 1st | 0 | 8 | 1 | 1 | 40 | 0 | 0.2 |
| | | | 2nd | 142 | 6 | | 0 | 2.5 | 0 | 0.5 |
| | | | 3rd | 163 | 8 | | 1 | 5 | 0 | 0.4 |
| | | | 4th | 233 | 10 | | 1 | 0 | 0 | 0.1 |
| S107 | 32 | White Hispanic | 1st | 0 | 14 | 1 | 1 | 20 | 0 | 1.0 |
| | | | 2nd | 38 | 22 | | 1 | 25 | 0 | 3.1 |
| S120 | 37 | African American | 1st | 0 | 15 | 1 | 1 | 40 | 0 | 3.9 |
| | | | 2nd | 215 | 2 | | 0 | 0 | 0 | 0.0 |
| S124 | 35 | White Hispanic | 1st | 0 | 16 | 1 | 1 | 20 | 0 | 0.0 |
| | | | 2nd | 114 | 8 | | 0 | 0 | 0 | 0.0 |
| | | | 3rd | 228 | 10 | | 0 | 5 | 1 | 0.0 |
| S134 | 33 | White | 1st | 0 | 4 | 1 | 0 | 0 | 0 | 0.0 |
| | | | 2nd | 201 | 6 | | 0 | 0 | 0 | 0.0 |
| | | | 3rd | 236 | 6 | | 0 | 0 | 0 | 0.0 |
| | | | 4th | 260 | 6 | | 0 | 0 | 0 | 0.0 |
| S172 | 24 | Asian | 1st | 0 | 12 | 1 | 1 | 0 | 0 | 0.0 |
| | | | 2nd | 34 | 9 | | 0 | 0 | 1 | 0.0 |
| | | | 3rd | 49 | 3 | | 0 | 0 | 0 | 0.0 |
| S202 | 38 | White | 1st | 0 | 16 | 1 | 1 | 60 | 1 | 0.0 |
| | | | 2nd | 57 | 2 | | 0 | 0 | 1 | 0.0 |

**Supplementary Table 1B. Demographic, clinical and treatment features of Lupus patients without renal involvement evaluated overtime.**

| Patient ID | Age | Ethnicity | Sample | Collection week span | SLEDAI | Renal ACR | Renal active | MEDs | | R. *gnavus* % |
|---|---|---|---|---|---|---|---|---|---|---|
| | | | | | | | | Prednisone | MMF | |
| S49 | 52 | African American | 1st | 0 | 2 | 0 | 0 | 0 | 0 | 0.4 |
| | | | 2nd | 41 | 3 | | 0 | 0 | 0 | 0.1 |
| S61 | 42 | Asian | 1st | 0 | 8 | 0 | 0 | 60 | 0 | 0.6 |
| | | | 2nd | 27 | 11 | | 0 | 3.75 | 0 | 4.7 |
| S188 | 57 | Asian | 1st | 0 | 9 | 0 | un | 0 | 0 | 0.1 |
| | | | 2nd | 24 | 5 | | un | 2.5 | 0 | 0.4 |
| S190 | 34 | White Hispanic | 1st | 0 | 4 | 0 | 0 | 0 | 0 | 0.0 |
| | | | 2nd | 67 | 4 | | un | 0 | 0 | 0.0 |
| S191 | 43 | White Hispanic | 1st | 0 | 4 | 0 | 0 | 0 | 0 | 0.0 |
| | | | 2nd | 54 | 0 | | 0 | 0 | 0 | 0.0 |
| | | | 3rd | 83 | 4 | | 0 | 0 | 1 | 0.2 |
| S198 | 35 | White Hispanic | 1st | 0 | 6 | 0 | 0 | 0 | 1 | 0.0 |
| | | | 2nd | 54 | 0 | | 0 | 0 | 1 | 0.0 |
| S205 | 46 | Asian | 1st | 0 | 4 | 0 | 0 | 0 | 0 | 0.0 |
| | | | 2nd | 31 | 6 | | 0 | 0 | 0 | 0.1 |
| | | | 3rd | 52 | 4 | | 0 | 0 | 0 | 0.0 |

**Supplementary Table 2A. SLEDAI domain scoring in patients with Lupus nephritis.**

| Patient ID | Sample | SLEDAI | Involvements in the SLEDAI Score |
|---|---|---|---|
| S47 | 1st | 8 | proteinuria, ↓complement, ↑dsDNA |
| | 2nd | 8 | proteinuria, ↓complement, ↑dsDNA |
| S78 | 1st | 8 | rash, alopecia, ↓complement,↑dsDNA |
| | 2nd | 6 | rash, pericarditis, ↑dsDNA |
| | 3rd | 4 | rash, ↑dsDNA |
| | 4th | 5 | rash, ↓WBC, ↑dsDNA |
| | 5th | 4 | rash, ↑dsDNA |
| | 6th | 8 | proteinuria, ↓complement, ↑dsDNA |
| S89 | 1st | 8 | proteinuria, ↓complement, ↑dsDNA |
| | 2nd | 6 | pleurisy, ↓complement, ↑dsDNA |
| | 3rd | 8 | proteinuria, ↓complement, ↑dsDNA |
| | 4th | 10 | proteinuria, pleurisy, ↓complement, ↑dsDNA |
| S107 | 1st | 15 | proteinuria, rash, alopecia, ulcers, ↓complement, ↓WBC, ↑dsDNA |
| | 2nd | 23 | arthritis, proteinuria, pyuria, rash, alopecia, pleurisy, ↓complement, ↓WBC, ↑dsDNA |
| S120 | 1st | 15 | myositis, proteinuria, pleurisy,↓ complement,↑dsDNA, fever |
| | 2nd | 2 | ↑dsDNA |
| S124 | 1st | 16 | visual disturbances, arthritis, ↓complement,↑dsDNA |
| | 2nd | 8 | arthritis, ↓complement, ↑dsDNA |
| | 3rd | 10 | arthritis, pericarditis, ↓complement, ↑dsDNA |
| S134 | 1st | 4 | ↓complement, ↑dsDNA |
| | 2nd | 6 | rash, ↓complement, ↑dsDNA |
| | 3rd | 6 | rash, ↓complement, ↑dsDNA |
| | 4th | 6 | rash, alopecia, ↓complement |
| S172 | 1st | 12 | hematuria, proteinuria, ↓complement, ↑dsDNA |
| | 2nd | 9 | proteinuria, ↓complement, ↓WBC, ↑dsDNA |
| | 3rd | 7 | proteinuria, ↓WBC, ↑dsDNA |
| S202 | 1st | 16 | hematuria, proteinuria, pyuria, ↓complement, ↑dsDNA |
| | 2nd | 2 | ↑dsDNA |

**Supplementary Table 2B. Organ involvements reflected by SLEDAI in non-Lupus nephritis patients**

| Patient ID | Sample | SLEDAI | Involvements in the SLEDAI Score |
|---|---|---|---|
| S49 | 1st | 2 | ↓complement |
| | 2nd | 3 | ↓complement , ↓WBC |
| S61 | 1st | 8 | rash, alopecia, ↓complement, ↑dsDNA |
| | 2nd | 11 | arthritis, alopecia, ↓complement, ↓WBC, ↑dsDNA |
| S188 | 1st | 9 | arthritis, ↓complement, ↓WBC, ↑dsDNA |
| | 2nd | 5 | ↓complement, ↓WBC, ↑dsDNA |
| S190 | 1st | 4 | ↓complement, ↑dsDNA |
| | 2nd | 4 | ↓complement, ↑dsDNA |
| S191 | 1st | 4 | arthritis |
| | 2nd | 0 | – |
| | 3rd | 4 | arthritis |
| S198 | 1st | 6 | arthritis, ↑dsDNA |
| | 2nd | 0 | – |
| S205 | 1st | 4 | ↓complement, ↑dsDNA |
| | 2nd | 6 | ↓complement, ↑dsDNA, alopecia |
| | 3rd | 4 | ↓complement, ↑dsDNA |

**Supplementary Table 3A. Other medications** of patients with Lupus nephritis

| Patient ID | Sample | Other medications |
|---|---|---|
| S47 | 1st | hydroxychlorquine 400 mg, azathioprine 150 mg |
| | 2nd | hydroxychlorquine 400 mg |
| S78 | 1st | hydroxychlorquine 400 mg |
| | 2nd | hydroxychlorquine 400 mg, milatuzumab/placebo trial |
| | 3rd | hydroxychlorquine 400 mg, methotrexate 20 mg |
| | 4th | hydroxychlorquine 400 mg, methotrexate 20 mg |
| | 5th | hydroxychlorquine 400 mg, methotrexate 20 mg |
| | 6th | hydroxychlorquine 400 mg, methotrexate 20 mg |
| S89 | 1st | hydroxychlorquine 400 mg |
| | 2nd | hydroxychlorquine 400 mg, belimumab 10 mg, azathioprine 100 mg |
| | 3rd | hydroxychlorquine 400 mg, belimumab 10 mg, azathioprine 100 mg |
| | 4th | hydroxychlorquine 400 mg, methylprednisolone 1000 mg |
| S107 | 1st | hydroxychlorquine 400 mg, anifrolumab/placebo study |
| | 2nd | hydroxychlorquine 400 mg |
| S120 | 1st | hydroxychlorquine 400 mg |
| | 2nd | hydroxychlorquine 200 mg, azathioprine 100 mg |
| S124 | 1st | hydroxychlorquine 400 mg |
| | 2nd | hydroxychlorquine 400 mg |
| | 3rd | hydroxychlorquine 400 mg |
| S134 | 1st | hydroxychlorquine 400 mg |
| | 2nd | hydroxychlorquine 300 mg, belimumab 10 mg |
| | 3rd | hydroxychlorquine 300 mg, belimumab10 mg |
| | 4th | hydroxychlorquine 300 mg |
| S172 | 1st | hydroxychlorquine 400 mg |
| | 2nd | hydroxychlorquine 400 mg |
| | 3rd | hydroxychlorquine 300 mg |
| S202 | 1st | hydroxychlorquine 400 mg, methotrexate 12.5 mg |
| | 2nd | hydroxychlorquine 300 mg |

**Supplementary Table 3B. Medications of patients without Lupus Nephritis**

| Patient ID | Sample | Other medications |
|---|---|---|
| S49 | 1st | hydroxychlorquine 400 mg, azathioprine 100mg |
| | 2nd | hydroxychlorquine 400 mg, azathioprine 150mg |
| S61 | 1st | hydroxychlorquine 400 mg |
| | 2nd | hydroxychlorquine 400 mg |
| S188 | 1st | hydroxychlorquine 400 mg |
| | 2nd | hydroxychlorquine 400 mg |
| S190 | 1st | hydroxychlorquine 400 mg |
| | 2nd | hydroxychlorquine 300 mg |
| S191 | 1st | hydroxychlorquine 400 mg |
| | 2nd | hydroxychlorquine 200 mg |
| | 3rd | hydroxychlorquine 200 mg |
| S198 | 1st | hydroxychlorquine 400 mg |
| | 2nd | hydroxychlorquine 400 mg |
| S205 | 1st | hydroxychlorquine 300 mg |
| | 2nd | hydroxychlorquine 400 mg |
| | 3rd | hydroxychlorquine 300 mg |
